# Single cell functional immunogenomics of the fallopian tube reveals a precursor immune surveillance network for ovarian cancer prevention

**DOI:** 10.1101/2025.06.30.662408

**Authors:** Luyao Wang, Breeshey Roskams-Hieter, Nosheen Hussain, Joel Nulsen, Aneesh Aggarwal, May Sallam, Lili Wang, Lena Rai, Amro Ahmed-Ebbiary, Aws Al-deka, Takashi Takeda, Mara Artibani, Hooman Soleymani majd, Jason Yap, Christopher Yau, Nancy Zaarour, Ahmed Ashour Ahmed

## Abstract

The fallopian tube (FT) is increasingly recognized as the site of origin for high-grade serous ovarian cancer (HGSOC), yet its immune landscape and potential role in tumor surveillance remain poorly understood. Here, we employ single-cell RNA sequencing (scRNA-seq) and paired single-cell T-cell receptor sequencing (scTCR-seq) to profile tissue-resident memory-like T cells (TRMLs) from matched non-cancerous FT, metastatic omental tumors, and peripheral blood in HGSOC patients. Surprisingly, we identify a substantial clonal and functional overlap between TRMLs from non-cancerous FT and tumor-infiltrating TRMLs, with 18.4% of tumor TRML clonotypes being shared with FT TRMLs, significantly exceeding overlap with circulating T cells. Shared TCR clonotypes are preferentially enriched in exhausted CD8+ T cells (CD8-Tex) and proliferative exhausted CD8+ subsets, suggesting prior antigenic exposure. Notably, we identify a previously uncharacterized CD8+ SIK3-high subset enriched in tumors, with gene expression signatures implicating epigenetic plasticity and metabolic adaptation.

Functionally, non-cancerous FT-derived TRMLs recognize autologous tumor antigens, as evidenced by robust IFN-γ responses in tumor organoid co-culture assays and CD137 upregulation. Whole-genome and transcriptomic analyses reveal shared somatic mutations between the FT and tumor, supporting the FT as the tumor’s evolutionary origin. Predicted tumor neoantigens elicit TRML responses in the non-cancerous FT, and single-cell TCR profiling confirms that neoantigen-reactive T cells in FT share clonotypes with tumor-infiltrating TRMLs, further reinforcing their role in early immune surveillance. Importantly, FT-derived TRMLs exhibit lower exhaustion signatures than their tumor counterparts, suggesting their potential as a preferable source for adoptive T cell therapies. Our findings uncover a precursor immune surveillance network in the FT and provide a rationale for leveraging FT-resident T cells in cancer immunotherapy and prevention.

## INTRODUCTION

Despite strong evidence linking T-cell infiltration to improved survival, ovarian cancer remains largely refractory to immunotherapy^1^, underscoring a critical knowledge gap in our understanding of the immune landscape at early disease stages. Therefore, a comprehensive understanding of the evolution of T cell responses against ovarian cancer, especially in the fallopian tube (FT) where high-grade serous ovarian cancer (HGSOC) arises^2–5^, would help in the development of novel immune-preventive strategies for ovarian cancer. Recent evidence suggests that the precancerous lesions including serous tubal intraepithelial carcinomas (STICs) occurring in the FT coincide with progressive shifts in the tumor microenvironment, transitioning from active immune surveillance, characterized by the presence of immune cells including tissue-resident memory (TRM) CD8+ T cells in early STICs to immune suppression in advanced STICs and cancer.^6–12^

HGSOC arises from the secretory cells of the FT epithelium.^13,14^ However, despite the aggressive metastatic nature of HGSOC, the primary FT lesions are confined to microscopic intraepithelial carcinomas, the so called serous tubal intraepithelial carcinomas (STICs). In contrast, abdominal metastases are capable of large macroscopic mass formation, fibrosis induction and distortion (e.g. omental cakes). The mechanism underlying the successful confinement in the FT remains unclear. We hypothesize that T cells in the FT play an active role in tumor immunosurveillance, potentially delaying or preventing ovarian cancer progression. However, the molecular and functional characteristics of these T cells remain undefined. Recent reports suggest that local immunity plays a critical role in maintaining a cancer-immune equilibrium.^15,16^ The reason this equilibrium is private to the FT but fails elsewhere in the abdominal cavity is an outstanding question to be answered. Delineating the molecular immune surveillance responses against early cancer development in the FT is indispensable for understanding the dynamic changes across the FT and metastatic ovarian cancer.

Recently, a unique subset of T cells, characterized by CD103+ expression, known as tissue-resident memory (TRM) T cells were found to play an important role in inhibiting cancer progression and metastasis, particularly in cancers of epithelial origin.^17–21^ Traditionally, TRM T cells are thought to differentiate from effector memory T cells and are primarily recruited into peripheral tissues in response to local inflammation but remain resident within the tissues long after the inflammation is resolved.^22–24^ The binding of CD103 to E-cadherin, which is expressed on epithelial cells, allows TRM cells to be maintained in the local tissue microenvironment and is believed to play critical roles in local immunosurveillance.^25^ TRM cells may be involved in cancer immunity from the time premalignant cells develop by residing in epithelial parts of peripheral tissues.^19^ Recently, TRM CD8+ T cells have been shown to contribute to shaping immunogenicity of ovarian cancer.^20^ However, their role in suppressing early ovarian cancer remains unclear. We hypothesize that TRMs might play an active role in local immunosurveillance by contributing to the successful confinement of STICs in the FT epithelium. Understanding the molecular features underlying this successful confinement requires paired analyses of TRMs from the apparent non-cancerous FT and metastatic tumors from the same patient. Such a comparison could enable the identification of key gene expression differences and enable hypothesis generation regarding the mechanistic basis of the apparent superiority of FT TRMs in confining preneoplastic lesions.

In this study, we leverage single-cell RNA sequencing (scRNA-seq) and paired single-cell TCR sequencing (scTCR-seq) to comprehensively characterize TRM-like cells (TRMLs) in FT and their relationship to ovarian tumor-infiltrating TRMLs. We complement this work with the use of organoids and immune-organoid co-culture systems to elucidate the functional responses of FT TRMLs to tumor antigens. We demonstrate that TRMLs from the FT share TCR clonotypes with tumor TRMLs and recognize tumor neoantigens, suggesting an intrinsic immune surveillance function that may be leveraged for immunotherapy and prevention. By elucidating the immune surveillance functions of FT TRMLs, this study provides a foundation for leveraging TRM biology in ovarian cancer prevention and immunotherapy. Given their less exhausted phenotype and tumor-reactive potential, FT-derived TRMLs could represent a promising source for adoptive T cell therapies. Additionally, the identification of tumor-reactive TCR clonotypes in FT raises the possibility of targeting these T cells in personalized neoantigen-directed immunotherapies, with implications for both therapeutic and preventive strategies in ovarian cancer.

## RESULTS

### Characterization of T cells by multisite scRNA-seq from HGSOC patients

To characterize the clonotypes of TRMLs in FT and TRM-like tumor-infiltrating lymphocytes (TILs) in HGSOC, we employed droplet-based 5`′` scRNA-seq and paired scTCR-seq. We profiled matched samples of CD3+CD103+ TRM-like cells (TRMLs) from non-cancerous FT tissues, paired Omentum tumor metastases and CD3+ T cells from peripheral blood mononuclear cells (PBMCs) from the same patients. Given the paucity of CD103+ cells in PBMCs, we opted to profile all CD3+ cells to enable clonotype tracking. We confirmed that the FTs were non-cancerous based on histopathology examination and next generation DNA sequencing (see later). Cells were sorted by fluorescence-activated cell sorting (FACS) from 4 different HGSOC patients (Figure 1A, Figure S1, Figure S2A). FACS analysis revealed that the mean percentage CD3+CD103+ TRMLs out of all CD3+ T cells in the FT was 8 times higher than TRM-like TILs (80 % and 11.4 %, respectively, Figure 1B-C). After quality-control filtering of scRNA seq data (see Methods) a total of 51,855 T cells remained (Figure 1D) of which 53.4% produced TCR α- and β-chain pairs. These T cells were further grouped into 12,206 distinct clonotypes by matching both α- and β-chain pairs, allowing us to track clonal lineages and measure clonal expansion in different tissue sites.

**Figure 1.**
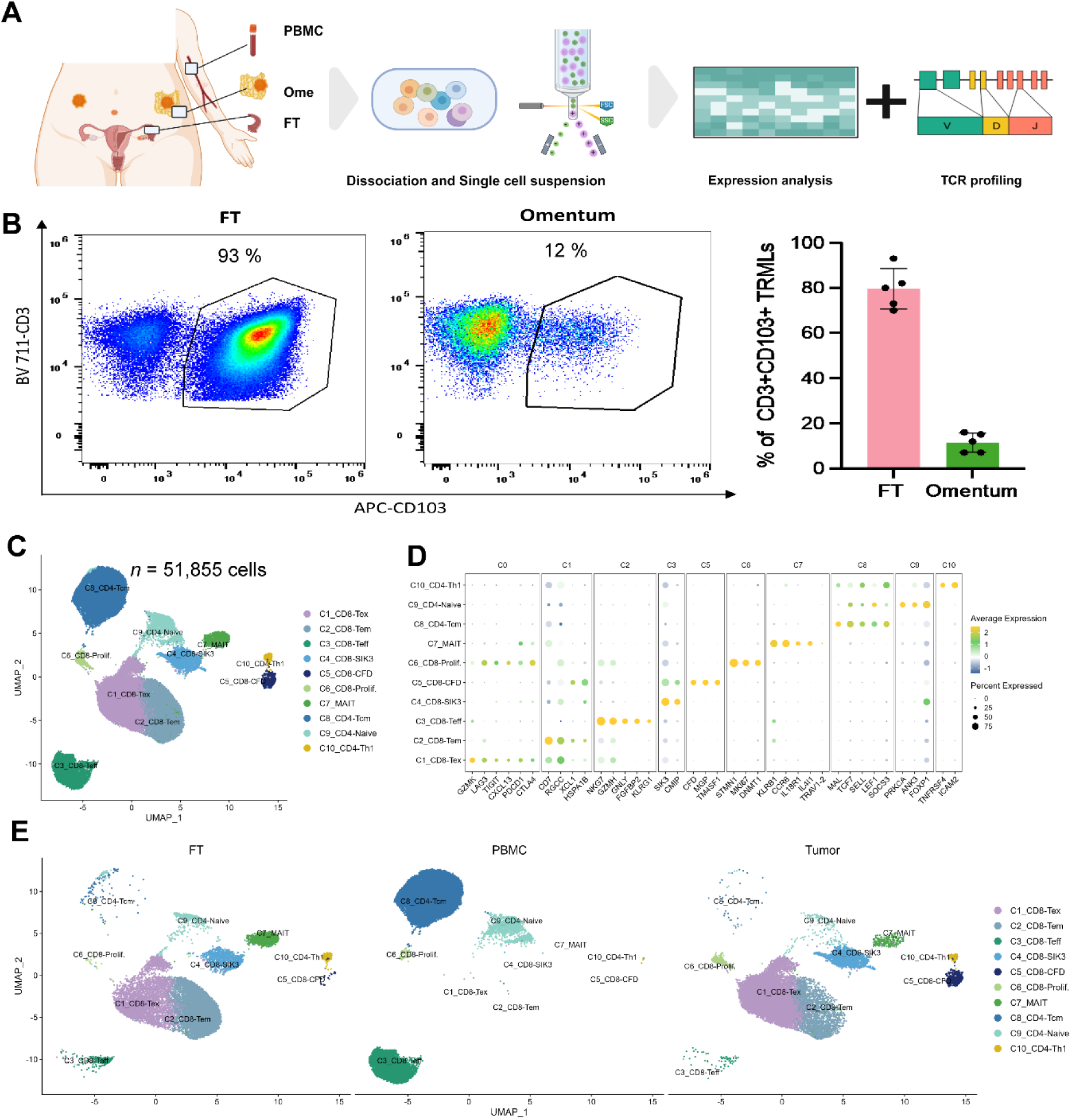
Landscape of immune cells of HGSOC via scRNA-seq of three sites. (A) Schematic representation of the study design, illustrating the workflow for sample collection, processing, and single-cell immune profiling in HGSOC patients using 10x Genomics sequencing. Samples were obtained from n = 4 patients, with matched Fallopian Tube (FT), Peripheral Blood Mononuclear Cells (PBMC), and Omentum tumor (Ome) tissues analyzed. (B) Representative dot plot showing co-expression of CD3 and CD103 on tissue-resident memory-like T cells (TRMLs) from paired FT and tumor samples, as assessed by flow cytometry. (C) Bar graph quantifying the frequency of CD3+CD103+ TRMLs as a proportion of total CD3+ T cells across biological replicates (n = 5). Data are presented as mean ± s.d. (D) Uniform Manifold Approximation and Projection (UMAP) visualization of single-cell transcriptomic data, identifying 10 distinct T cell clusters across n = 4 HGSOC patients. Each dot represents a single cell, color-coded by cluster identity: C1_CD8-Tex: CD8+ Exhausted T Cells, C2_CD8-Tem: CD8+ Effector Memory T Cells, C3_CD8-Teff: CD8+ Effector T Cells, C4_CD8-SIK3: CD8+ SIK3-high T Cells, C5_CD8-CFD: CD8+ CFD-high T Cells, C6_CD8-Prolif.: CD8+ Proliferating T Cells, C7_MAIT: Mucosal-Associated Invariant T Cells, C8_CD4-Tcm: CD4+ Central Memory T Cells, C9_CD4-Tnaive: CD4+ Naïve T Cells, C10_CD4-Th1: CD4+ Th1 T Cells. (E) Dotmap displaying the expression levels of key marker genes across major T cell clusters in HGSOC samples. Rows represent individual clusters, and columns represent selected genes. (F) UMAP plots illustrating distinct immune cell compositions across the three analyzed tissue sites (FT, PBMC, and Tumor) in HGSOC patients, highlighting site-specific immune profiles.

We identified 10 clusters; 6 CD8+ subsets, 3 CD4+ subsets, and one cluster of MAIT T cells (Figure 1D). Of these 6 CD8+ subsets, three were characterized by expression of activation markers *GZMK*, *GZMB*, and exhaustion markers *LAG3*, *TIGIT*, *PDCD1*, *CTLA4*, and chemokine *CXCL13*, associated with B cell recruitment and neoantigen burden in human malignancies(Figure 1D, E and S2B-C).^26^ Differential gene expression analyses revealed that one of these clusters (C1_CD8-Tex) expressed particularly high levels of exhaustion markers, suggestive of prolonged exposure to cognate antigens; while another (C6-CD8- Prolif.) cluster, in addition to the aforementioned markers, expressed genes associated with cell cycle progression and proliferation (e.g., *MKI67*, *DNMT1*, and *STMN1*). The third (C2_CD8-Tem) was distinguished by the expression of genes that are typically induced after activation (e.g., *RGCC*, *XCL1*, and *HSPA1B*). One other cluster(C3_CD8-Teff) stood out based on the expression of genes associated with cytotoxic activity and its presumed terminally differentiated nature (e.g., *NKG7*, *GZMH*, *GNLY*, *FGFBP2*, *KLRG1*). C4_CD8-SIK3 appeared unconventional with high expression of *SIK3* (Salt-Inducible Kinase 3), a member of the AMPK family of kinases. Another cluster (C5_CD8-CFD) displayed high *CFD* expression, a key regulator of the alternative complement pathway that links innate and adaptive immunity, influencing T cell function and immune regulation.^27^

Within the CD4+ T cell compartment, we detected three clusters (Figure 1D-F and S2B-C). We observed 2 CD4+ clusters that appeared highly enriched in PBMCs that expressed stemness markers (*TCF7*, and *SELL*). One of them, C8_CD4-Tcm, expressed high levels of the central memory marker gene *MAL*, and the other one (C9_CD4-Tnaive) highly expressed *PRKCA*, *ANK3*, and *FOXP1*, compatible with naïve T cell state. C10_CD4-Th1 cluster was distinguished by expression of T helper cell markers *TNFRSF4*, and *ICAM2*. Finally, we observed a cluster (C7_MAIT) marked by high expression of genes typically associated with Mucosal-Associated Invariant T cells (*KLRB1*, *CCR6*, *IL18R1*, *IL4I1*, *TRAV1-2*).^28–31^

Notably, we found the TRMLs from FT and tumor sites exhibit similar cellular compositions (Figure 1F, and Figure S2A, D). However, the C1_CD8-Tex cluster (Cluster 1, CD8+ exhausted T cells) represented 61% of T cells in the tumor sample compared with 25.4% in FT (Figure S2E). In contrast, the C2_CD8-Tem cluster (Cluster 2, CD8+ effector memory T cells) was the most prominent in FT samples, accounting for 50% of T cells in the FT sample compared with 14.9% in the tumor. Additionally, cells within the C6_CD8- Prolif. cluster (Cluster 6, CD8+ proliferating T cells) were more abundant in tumor samples compared to FT-derived TRMLs (5.8% and 1.2%, respectively, Figure S2E). As expected, the T cells from the PBMC samples displayed a markedly different cellular composition compared to those from FT or tumor samples, with the most abundant population being the C8_CD4-Tcm cluster (Cluster 8, CD4+ central memory T cells, 60.2 %) and the C3_CD8-Teff cluster (Cluster 3, CD8+ effector T cells, 29.1%), which were rarely observed in FT and tumor samples. All three sample types shared the C6_CD8-Prolif. cluster (Cluster 6, CD8+ proliferating T cells) (Figure S2E). C5_CD8-CFD cluster that was characterized by high expression of *CFD* and C10_CD4-Th1 cluster that was characterized by expression of T helper cell markers *TNFRSF4, and ICAM2* were patient specific clusters, all other cells contained cells from multiple patients (Figure S2F), cluster 5 and cluster 10 were, therefore, not considered for further detailed analysis.

### TCRs from FT were highly shared with TCRs from Tumor

Previous reports using immunohistochemistry documented the presence of immune infiltrates around precancerous lesions in the fallopian tube in patients with HGSOC.^6,32^ We first reasoned that if apparent non-cancerous FTs were the site of origin of metastatic HGSOC, then their T cells would have developed a memory for the tumors that were once contained within the FT. Since TCRs provide a reliable molecular tag to track the lineage of T cells, we first looked at the overall clonal diversity across different tissues and found the PBMC samples showed the highest unique clones (8,895 detected TCRs) compared to T cells from FT and tumor, which showed a similar number of unique clones with each other (2,415 TCRs in FT vs 2,029 TCRs in tumor) (Figure S3A). We further inspected unique clones individually across patients from the 3 different tissue sites, and found a similar trend to the aggregated analysis, with some fluctuation across patients within each tissue type (Figure S3B). We next compared the TCR-based clonotypes of TRMLs in the non-cancerous FTs and metastatic tumors from the same patients. Remarkably, we found that out of 2,029 tumor TRML TCR clonotypes from four patients, 373 (18.4%) were shared with those from FT, compared with only 180 (8.8%) shared with those from PBMCs (*P* = 2.2 × 10^−16^, Fisher’s Exact Test, Figure 2A). When analyzing individual patient samples, we observed that overlap rates ranged from 17.7% to 28.1% (Figure 2B), reflecting 43 to 199 clonotypes that were shared (Figure S3C), compared with an overlap of 0.7%-9.3% with PBMCs (Figure 2B), indicating a significantly higher degree of TCR overlap between FT and tumor.

**Figure 2.**
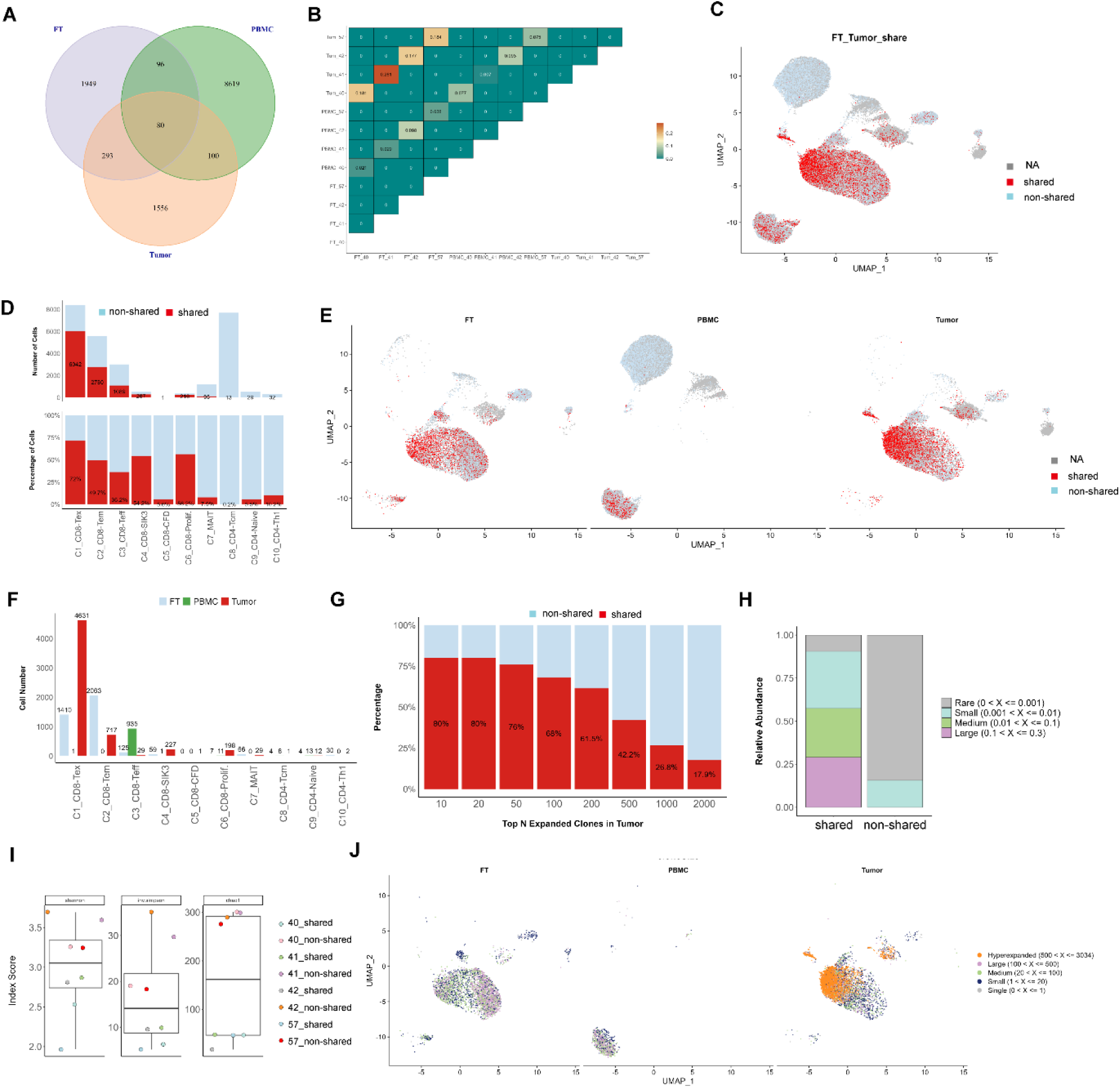
High degree of TCR overlap between fallopian tube (FT) and tumor-infiltrating T cells in HGSOC. The Venn diagram (A) presents the number of unique TCRs in FT (purple), tumor (orange), and PBMC (green), along with the overlap between these compartments. The clonal overlap plot (B) quantifies the percentage of shared TCRs across different tissue sites for each patient. UMAP visualization (C) highlights T cells with shared TCRs between tumor and FT (red), while non-shared TCRs are marked in light blue, and T cells lacking detected TCRs are in grey. The bar plots (D) illustrate the number and proportion of shared T cells across different clusters. Another UMAP plot (E) further delineates shared TCRs across distinct tissue sites. The bar graph (F) compares the number of shared TCRs within each cluster across FT (blue), PBMC (green), and tumor (red). The degree of TCR overlap between FT and the most expanded tumor clones is analyzed in (G), showing the percentage of shared TCRs from the top 10 to top 2000 expanded clones. The clonal abundance of shared versus non-shared TCRs is depicted in (H), categorizing T cell clones as rare (0 < X ≤ 0.001), small (0.001 < X ≤ 0.01), medium (0.01 < X ≤ 0.1), large (0.1 < X ≤ 0.3). Productive clonality analysis (I) compares the diversity of shared and non-shared TCRs using Shannon, Inverse Simpson, and Chao1 indices. The UMAP plot (J) shows the distribution of shared TCR clones across different tissue sites, classifying them based on clonal expansion: single (n = 1), small (1–20 cells), medium (20–100 cells), large (100–500 cells), and hyperexpanded (500–3034 cells). These results suggest that FT TRMLs exhibit significant clonal overlap with tumor-infiltrating T cells, reinforcing their potential role in early immune surveillance and tumor-reactive immunity.

Next, we integrated our scTCR-seq and scRNA-seq data to examine the distribution of shared TCR clonotypes. Notably, these shared clonotypes were predominantly enriched in the C1_CD8-Tex cluster. Of all TCRs that were contained within this cluster, 72% (n=6,042 cells) were shared TCRs, with only 28% of cells containing non-shared TCRs (*P* = 6.5× 10⁻¹^0^, Fisher exact test, Figure 2C-D). Furthermore, cells with shared TCRs were also enriched in the other CD8+ clusters; C2_CD8-Tem (2,780 cells, 49.7% of this cluster), C3_CD8-Teff (1,089 cells, 36.2% of this cluster), C4_CD8-SIK3 (287 cells, 54.2% of this cluster), and C6_CD8-Prolif. (216 cells, 56.2% of this cluster) (Figure 2C-D). These clusters are possibly highly related to tumor reactive T cells (TRMLs with exhausted or proliferating features).^33,34^ We next examined the abundance of cell cluster types in cells with shared TCRs in different tissues. We found that most cells with shared TCRs formed part of the tumor C1_CD8-Tex cluster, the FT C2_CD8-Tem and C1_CD8-Tex clusters and to a lesser extend the C3_CD8-Teff cluster in PBMC (Figure 2E-F). This observation suggests that these shared TCR clonotypes may represent tumor-reactive TCRs recirculating to the bloodstream, antimicrobial TCRs, or potentially a combination of both. However, the percentage of shared TCRs was particularly high among the top 10 and top 20 expanded clones, reaching 80%. Notably, the percentage overlap was directly proportional to the size of the tumor clone with the percentage dropping to only 17% when the least expanded clones were considered (Figure 2G). The link between tumor clonotype size and the percentage of cells with shared TCRs suggests that these cells are tumor-reactive.

Further clonality analysis revealed that cells with shared TCRs are more expanded than cells with non-shared TCRs (Figure 2H). More than 50% of clones exhibited high expansion rates (0.01∼0.3 expansion rate). In contrast, non-shared TCRs exhibited a lower expansion rate (expansion rate < 0.01, p-value < 2.2e-16, Pearson’s Chi-squared test). These results were consistent with the clonality diversity analysis, which revealed that non-shared cells displayed a higher Shannon Index (p-value = 0.009, t-test), Inverse Simpson Index (p-value = 0.027, t-test), and Chao1 Index (p-value = 1.885e-05, t-test) (Figure 2I). These results indicate that cells with shared TCRs were less diverse but more expanded, further suggesting that they were targeting specific tumor antigens.

We next examined the clonotype expansion and cluster distribution of cells with shared TCRs in different tissue types. Cells with shared TCRs from FT and PBMC were less expanded compared to cells from tumors (The clone proportions with expansion rates above 0.1 are 51.9% in tumors, 34.3% in PBMCs, and 11.7% in FT, Figure S3D). Notably, hyperexpanded T cells with shared TCRs (clonotypes > 500 cells and clonesize > 0.3 ) only appeared in the tumor and were mainly distributed in the C1_CD8-Tex, C4_CD8- SIK3, and C6_CD8-Prolif. Clusters (Figure 2J, S3D). These findings support that shared TCRs from FT were less expanded and less exhausted, with implications for cell immunotherapy.

Altogether, the above results support the notion that tumor clones with shared TCRs may be tumor-specific and that the FT clones with shared TCRs may be reactive to the same tumor antigens. However, the findings thus far do not exclude the possibility that these shared clonotypes might have developed due to prior exposure to common microbial infections. We further examined the TCR sharing between the tumor and PBMCs, used as a control, and observed a distinct pattern compared to the TCR overlap between FT and tumor. These shared clonotypes were predominantly enriched in the C3_CD8-Teff cluster within PBMC, comprising 1399 cells (46.6%) of the total T cells in this cluster, whereas in FT and tumor sites, the shared TCRs were most enriched in the C1_CD8-Tex and C2_CD8-Tem clusters (Figure S3E-G). Furthermore, we analyzed the percentage of shared TCRs between PBMC and tumor across the top 10 to 2000 most expanded clones in the tumor. Unlike the pattern observed with FT and tumor, the overlap ratio between PBMCs and tumor showed minimal variation, ranging from 7.6%∼30%, even as the number of top expanded clones in the tumor increased (Figure S3H). Considering the higher clonal diversity observed in PBMC compared to FT, but the lower overlap ratio with tumor, this pattern further supports the hypothesis that the shared TCRs between tumor and FT may be tumor-reactive. The overlap between PBMCs and tumor TCRs could be attributed to anti-microbial TCRs, tumor-reactive TCRs recirculating into the blood, or the possibility that the shared TCRs in PBMC represent progenitor TRMLs that are mobilized to combat the tumor.

### Cells with shared TCRs follow similar developmental paths but exist in distinct functional states in the FT and tumor

We next investigated how shared TCR-bearing cells transition between functional states by performing trajectory inference using Monocle3 (Figure 3A; STAR Methods). For this pseudotime analysis, we excluded MAIT cells due to their unique TCR characteristics, as well as CD8-Teff and CD4+ T cell subsets, which were largely restricted to PBMCs. The original CD8_Tex population was subdivided into two distinct clusters, CD8_Tex1 and CD8_Tex2. The inferred trajectory began with CD8_Tem cells, which first transitioned into CD8_Tex1. From there, the developmental pathway branched into three distinct endpoints: CD8_SIK3, CD8_Tex2, and CD8_Prolif (Figure 3A-B). When analyzing FT and tumor samples separately, we found that early-stage shared TCRs were predominantly present in FT samples, primarily following the trajectory CD8_Tem → CD8_Tex1 → CD8_SIK3. In contrast, shared TCRs in tumor samples were mostly positioned at terminal points of the exhaustion continuum, following paths such as CD8_Tex1 → CD8_SIK3, CD8_Tex2, or CD8_Prolif (Figure 3C).

**Figure 3.**
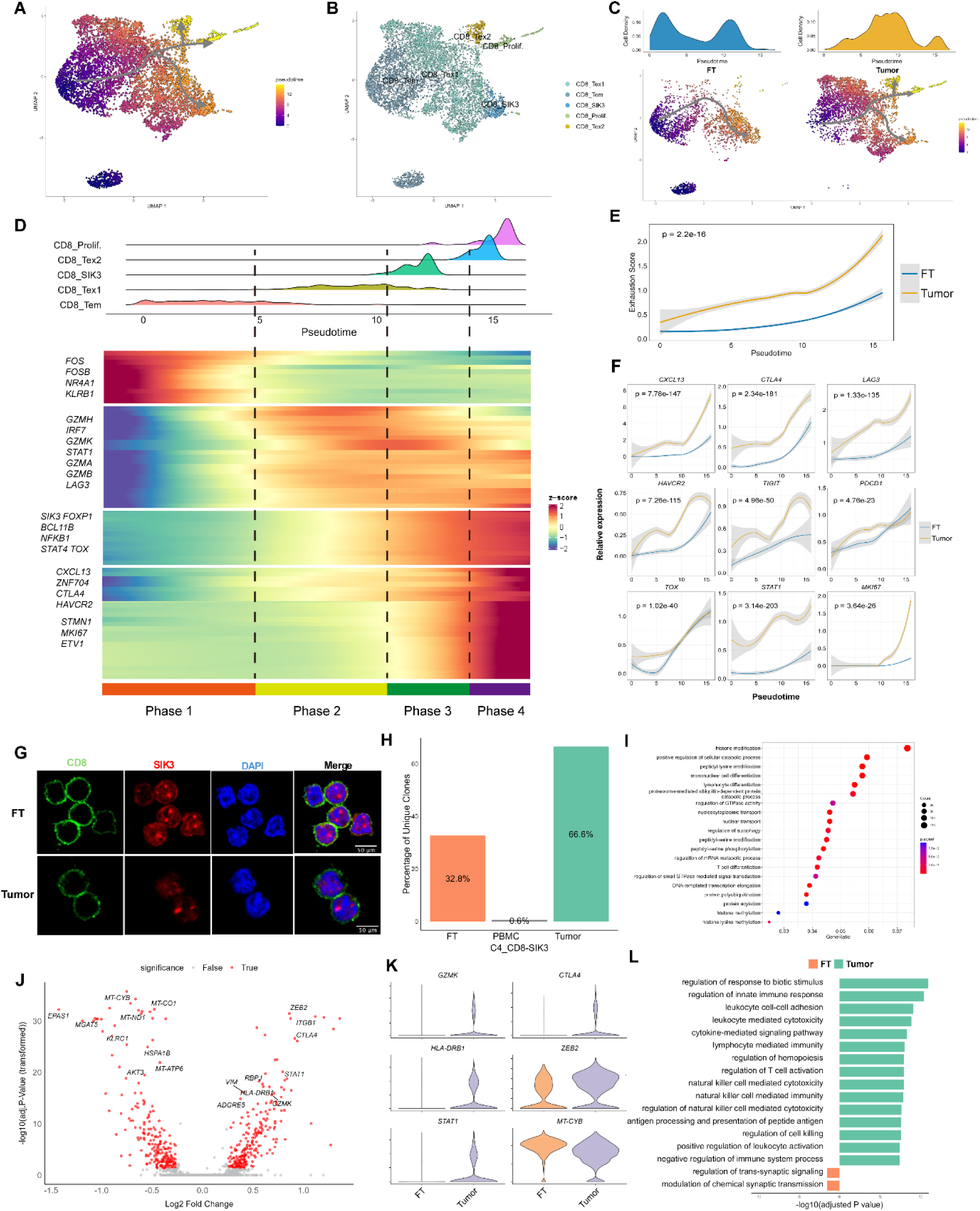
Transition trajectories of shared TCRs in FT and tumor samples, follow a similar trajectory, but occupy distinct transcriptional states. (A) Pseudotime trajectory mapping illustrates the differentiation paths of T cells bearing shared TCRs from fallopian tube (FT) and tumor samples. (B) T cell subtypes with shared TCRs are color-coded to indicate their distribution across the pseudotime continuum. (C) Shared TCR trajectories are displayed separately for FT-derived (left) and tumor-derived (right) samples, with cell state densities shown at the top of each panel. (D) Heatmap depicting dynamic gene expression changes along pseudotime (lower panel), with the upper panel showing the phase distribution of shared TCRs during transition, subdivided into four distinct phases and color-labeled by subtype. (E) Smoothed expression profiles of exhaustion scores across pseudotime reveal progressive changes; shaded gray bands represent 95% confidence intervals. (F) Pseudotime expression dynamics of selected marker genes, with Wilcoxon test significance and 95% confidence intervals indicated by error bars. (G) Immunofluorescence (IF) staining of SIK3 in CD8+ cells from HGSOC patient tissues (N=2) confirms co-localization of SIK3 (red) with CD8 (green), with nuclei stained by DAPI (blue). The robust nuclear staining of SIK3 in CD8+ cells mirrors transcriptional findings, suggesting a role for SIK3 in modulating CD8+ T cell function within both FT and tumor microenvironments. (H) A bar plot quantifies the percentage of unique clones in the C4_CD8-SIK3 cluster across FT, PBMC, and tumor samples, underscoring their differential presence in various tissue sites. (I) Differentially expressed genes activated pathways in the C4_CD8-SIK3 cluster compared to all other cells in the dataset are identified using Gene Ontology (GO) analysis, with an adjusted P value <0.01 (BH procedure). (J) A volcano plot highlights differentially expressed genes between the C4_CD8-SIK3 cluster in tumor versus FT, with genes meeting an adjusted P value threshold of <0.05 and log2(fold change) >0.25. (K) Violin plots display the expression levels of selected marker genes in FT and tumor-derived C4_CD8- SIK3 cells, emphasizing their differential activation states. (L) Additional pathway analysis distinguishes the transcriptional profiles of C4_CD8-SIK3 cells in tumor versus FT, identifying GO pathways with an adjusted P value <0.01.

We then examined the transcriptional dynamics accompanying the transitions of shared TCR-expressing cells and identified four distinct transcriptional phases along the trajectory (Figure 3D). Phase 1 was primarily composed of CD8_Tem cells, marked by elevated expression of AP-1 family transcription factors (*FOS*, *FOSB*), immediate early genes such as *NR4A1*^35^, the memory T cell marker *KLRB1*, and low levels of cytotoxic genes like *GZMA*, G*ZMB*, and *GZMH*—indicating a resting state with limited cytotoxic potential. Phase 2 cells displayed peak expression of classical cytotoxic genes (*GZMH*, *GZMB*, *GZMA*, *GZMK*, *IRF7*) alongside the early exhaustion marker *LAG3*, suggesting they represent a transitional, pre-exhausted state that maintains effector function while beginning to adopt exhaustion features due to persistent antigen exposure.

Phase 3 was notable for the upregulation of *SIK3*, a kinase involved in regulating cellular metabolism, including gluconeogenesis and lipid homeostasis. This phase also showed enriched expression of transcription factors such as *FOXP1*, *BCL11B*, *NFKB1*, and *TOX*. *FOXP1* is known to suppress proliferation and granzyme B (GZMB) expression in CD8+ T cells^36^, acting as a marker of functional restraint and contributing to the maintenance of memory T cell survival through TGF-β-mediated repression of c-JUN.^37^ These signatures suggest that Phase 3 represents a metabolically adapted and epigenetically modulated state with limited effector function.

Phase 4 was defined by elevated levels of terminal exhaustion markers, including transcription factors *ETV1* and *ZNF704*, and immune checkpoint genes such as *CTLA4*, *HAVCR2*, and chemokine *CXCL13*. *CXCL13*, in particular, has been previously shown to mark terminally exhausted CD8+ T cells specifically within the tumor microenvironment and is often upregulated in neoantigen-specific T cells, distinguishing them from non-reactive bystanders.^38–43^ This supports the notion that these cells are not only exhausted but also tumor-specific. Although proliferation markers like *MKI67* and *STMN1* were also upregulated in this phase, their co-expression with exhaustion markers suggests a dysfunctional proliferative state.

We further validated these patterns by demonstrating that exhaustion signatures progressively increased along pseudotime. Cells in the CD8_Tex2 and CD8_Prolif. clusters, which reside at the terminal ends of the trajectory, exhibited the highest exhaustion scores, indicating that these populations represent the endpoint of chronic antigen stimulation and functional exhaustion (Figure 3D-E).

To better understand the transitional states of shared TCR-expressing cells in the FT versus tumor tissue, we compared the key transcriptional changes associated with pseudotime progression in both sites. Across the trajectory, we observed a steady upregulation of exhaustion markers, including *CXCL13*, *CTLA4*, *LAG3*, *HAVCR2*, *TIGIT*, and *PDCD1*, as well as the exhaustion-related transcription factor *TOX*, the activation-related transcription factor *STAT1*, and the proliferation marker *MKI67* (Figure 3F). Importantly, all of these genes were expressed at significantly higher levels in tumor samples than in FT. At the terminal ends of the pseudotime trajectory, tumor-infiltrating T cells showed a pronounced increase in exhaustion markers— particularly *CXCL13* and *LAG3*—as well as *MKI67*, indicative of a hyperactivated yet dysfunctional state. In contrast, T cells from the FT demonstrated a more gradual rise in these markers, suggesting that they are less exhausted and potentially more functionally competent than their tumor counterparts.

Notably, our pseudotime analysis identified the CD8_SIK3-high cluster, which arises following CD8_Tex1 but does not exhibit features of terminal exhaustion, suggesting a distinct functional state. The presence of this branch in both FT and tumour suggests that CD8_SIK3-high represents an alternative fate within the exhaustion continuum. Given its unique trajectory and potential functional relevance, we further investigated the characteristics of this cluster.

The C4_CD8-SIK3 cluster is distinguished by high expression of Salt-Inducible Kinase 3 (*SIK3*), an enzyme known for its role in metabolic regulation, including processes like gluconeogenesis and lipid metabolism.^44^ We validated SIK3 expression at the protein level using immunofluorescence staining of CD8+ T cells from both tumor and FT tissues, where SIK3 was found to be broadly distributed throughout the cell, with a predominant nuclear localization likely corresponding to nucleosome-like structures (Figure 3G). While SIK3’s functions have been described in other cell types, its role in lymphocytes remains poorly understood. Notably, a recent study using a SIK3 knockout mouse model implicated this kinase in thymic T cell development.^45^ In our analysis, the SIK3-expressing CD8+ T cell subset was highly enriched in tumor tissue, comprising 67% of this cluster’s population, compared to 32.9% in the FT and minimal representation in PBMCs (Figure S2D). Moreover, clonal diversity within the C4_CD8-SIK3 cluster was markedly greater in tumors, accounting for 66.6% of the cluster’s clonotypes (Figure 3H). Pathway enrichment analysis showed strong associations with histone modification, cell catabolism, and differentiation processes (Figure 3I), pointing to an epigenetic regulatory function. These findings are consistent with prior studies that have linked SIK3 activity to histone deacetylase regulation in non-immune cells.^44,46^ The prominence of pathways involved in metabolic reprogramming suggests that C4_CD8-SIK3 cells may possess a high degree of epigenetic plasticity, enabling them to adapt to dynamic conditions in the tumor microenvironment such as hypoxia or chronic inflammation. Additionally, the enrichment of cell catabolic pathways implies a potential role in sustaining T cell homeostasis through metabolic adaptation.

Comparative gene expression analysis of the C4_CD8-SIK3 cluster between tumor and FT tissues revealed that in tumors, these cells exhibited upregulation of genes associated with both effector functions and T cell exhaustion, including *GZMK*, *CTLA4*, *HLA-DRB1*, *ZEB2*, and *STAT1* (Figure 3J-K). Conversely, there was notable downregulation of mitochondrial genes involved in oxidative phosphorylation, such as *MT- ND1*, *MT-CYB*, and *MT-ATP6* (Figure 3J-K). This suppression of mitochondrial activity likely reflects a shift away from oxidative phosphorylation toward glycolysis and lipid metabolism—a hallmark of metabolically reprogrammed immune cells responding to hypoxia and the high energy demands within the tumor microenvironment.^47–50^ This downregulation of mitochondrial genes was not unique to the SIK3 cluster in the tumor suggesting that the upregulation of SIK3 was more relevant to adaptation rather than steady state expression. Pathway enrichment analysis of the upregulated genes in tumor-derived C4_CD8-SIK3 cells showed strong associations with immune processes including T cell activation, leukocyte-mediated cytotoxicity, natural killer cell cytotoxicity, and cell-cell adhesion (Figure 3L). These features are consistent with a scenario in which C4_CD8-SIK3 cells in the tumor are actively responding to tumor antigens. In contrast, C4_CD8-SIK3 cells in the FT displayed a more quiescent phenotype, suggesting a homeostatic, surveillance-oriented function rather than direct effector engagement. Altogether, these findings support a model in which T cells bearing shared TCRs follow a similar developmental trajectory in both FT and tumor tissues but acquire distinct transcriptional and functional identities shaped by their local microenvironments.

### The TRM signature of shared TCRs between FT and the top 50 expanded T cell clonotypes from tumors may define neoantigen-reactive T cells

To determine whether the cells with shared TCRs might be tumor-reactive, we further performed differential gene expression analysis to characterize their transcriptomic signatures. We observed that cells with shared TCRs compared to non-shared TCRs differentially expressed 631 genes (Figure 4A). We found that the cells with shared TCRs highly expressed genes related to activation/effector genes (*GZMK*, *GZMA*, *CCL5*, *GZMB*, *HLA-DRA*, *HLA-DRB1*), TRM genes(*CXCL13*, *ZNF683*, *ITGAE*, *CCL3*), exhaustion-related genes (*CTLA4*, *PDCD1*, *HAVCR2*, *TIGIT*, *LAG3*), and proliferation-related genes(*STMN1*) (Figure 4B-C), which suggested that cells with shared TCRs were significantly more activated and exhausted compared to others irrespective of the tissue of origin lending further support to the hypothesis that these cells were tumor reactive.

**Figure 4.**
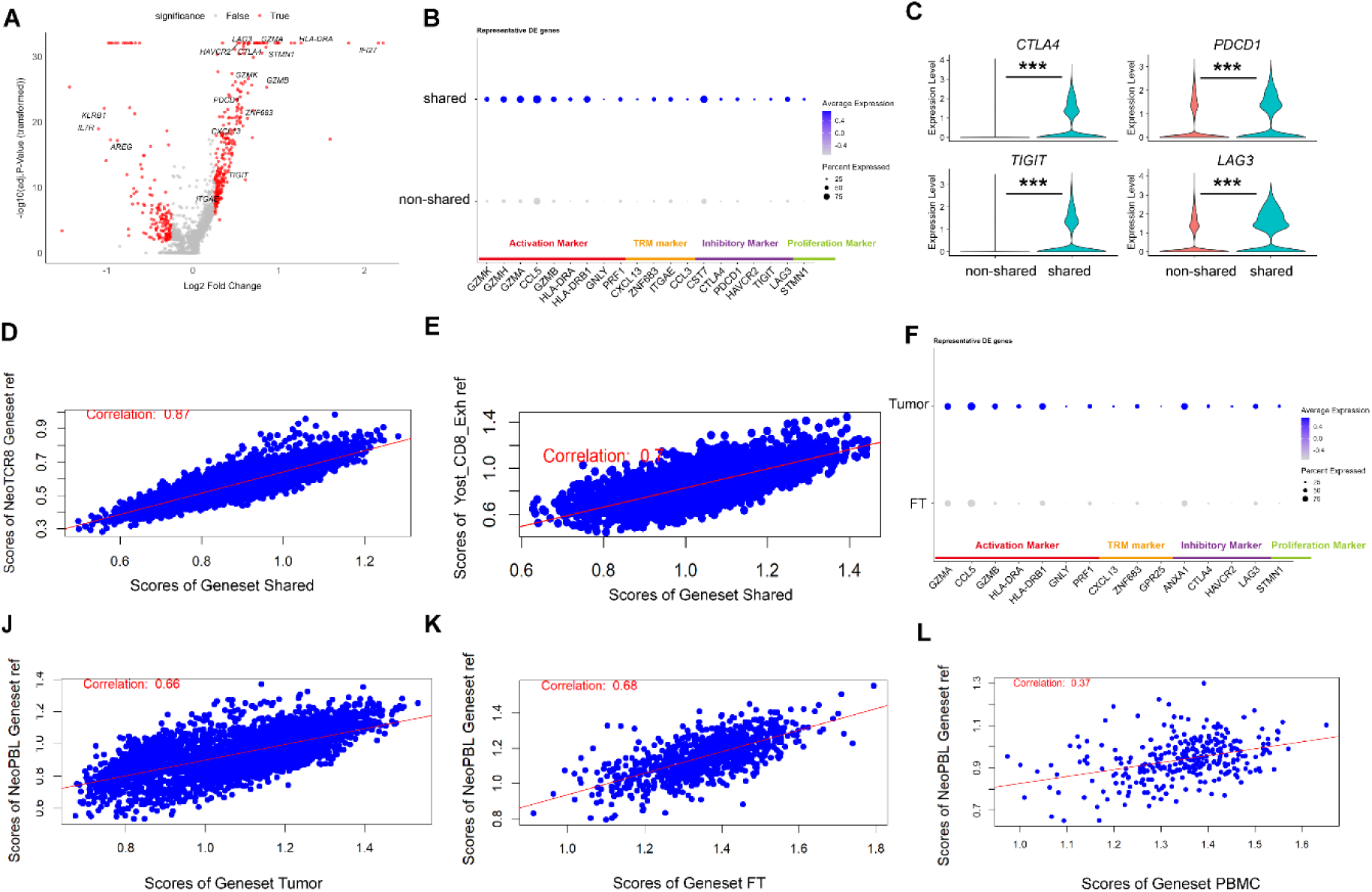
T cells with shared TCRs between FT and the top 50 expanded T cell clonotypes from tumors possess an exhaustion signature. (A) A volcano plot displays differentially expressed genes in shared TCR T cells versus non-shared TCR T cells, with red dots indicating genes meeting the statistical threshold of adjusted P < 0.05 and log2 fold change >0.25. (B) A dot plot highlights representative differentially expressed (DE) genes between shared and non-shared TCR T cells. (C) A violin plot illustrates normalized gene expression levels of key DE genes involved in T cell exhaustion, comparing shared and non-shared TCR T cells, with significance determined by a two-sided Wilcoxon test (***p<0.001). (D) A correlation plot evaluates the overlap between the DE geneset from shared versus non-shared TCR T cells and a previously published geneset of neoantigen-reactive CD8 T cells (NeoTCR8 Geneset ref.), with enrichment scores calculated via single-cell Gene Set Enrichment Analysis (scGSEA). (E) A correlation plot compares the DE geneset of shared versus non-shared TCR T cells against a known signature of CD8 exhausted T cells (Yost_CD8_Exh ref.), also using scGSEA-calculated enrichment scores. (F) A dot plot presents the DE genes distinguishing shared TCRs originating from tumors versus those from FT. (J-L) Additional correlation plots assess the relationship between DE genesets of shared versus non-shared TCR T cells and a published geneset of neoantigen-reactive CD8 T cells from PBMC (NeoPBL Geneset ref), separately analyzed within tumor (J), FT (K), and PBMC (L) datasets.

Next, we assessed the correlation between our candidate gene set (geneset ‘Shared’ that comprises 365 upregulated genes that showed an adjusted p_value of less than 0.05 and absolute log2FC of more than 0.25) and the validated and published NeoTCRs gene set.^51^ Utilizing Gene set variation analysis (GSVA) to calculate the scGSEA score, we found that geneset ‘Shared’ demonstrated a strong correlation (*r* = 0.87) with the NeoTCR8 geneset (Figure 4D).

Furthermore, our geneset strongly correlated (*r* = 0.7) with the published dysfunctional gene set^52^ suggesting that cells with shared TCRs exhibit an exhausted phenotype but are not restricted to typical exhaustion markers (Figure 4E). Previous studies have identified neoantigen-reactive TCRs as tumor-responsive, with the corresponding T cells displaying mixed transcriptional signatures beyond classical exhaustion markers.^51,53^ These findings imply that the geneset ‘Shared’ signature identified in HGSOC could serve as a marker of tumor-reactive T cells. However, further studies are necessary to validate whether this signature can effectively distinguish between bystander microbial-reactive TILs and those reactive to tumor-associated viral antigens.

Since T cells within the tumor are, presumably, under sustained stimulation by tumor antigens, we reasoned that they are potentially more exhausted than their corresponding clones in the FT. To test this, we performed differential gene expression analysis between T cells with shared TCRs in the FT and their counterparts in the tumor (Figure 4F). This analysis revealed that key exhaustion markers such as *CTLA4*, *HAVER2*, *LAG3* and activation markers such as *GZMA*, *GZMB*, *HLA-DRA*, *HLA-DRB1* were profoundly underexpressed in the FT clones compared to their respective clones in the tumor (adjusted p-value of less than 0.05 and absolute log2FC of more than 0.25, Figure 4F) This observation suggests that FT-derived T cells with shared TCRs, are much less exhausted and could, therefore, be promising candidates for adoptive T cell therapy.

We next compared T cells with shared TCRs to non-shared TCRs in the tumor, FT and PBMC independently (Figure S4A-C). We found that cells with shared TCRs were significantly more activated and exhausted compared to non-shared cells irrespective of the tissue of origin (Figure S4A-C). Pathway enrichment analysis (Figure S4D-F) revealed an enrichment of pathways involved in T cell differentiation, activation, and adhesion, suggesting that cells with shared TCRs appear to be dynamically stimulated presumably by tumor antigens. Interestingly, this analysis identified enrichment of viral response pathways only in the tumor site and not in the FT TRMLs, raising the possibility that persistent T cell activation by tumors elicits pathways that are similar to those activated by anti-viral immunity.

We next examined how the transcriptional signatures of shared TCRs correlated with established neoantigen-reactive T cell profiles based on their tissue of origin. When comparing the tumor-derived gene set (316 genes upregulated in shared versus non-shared TCRs within the tumor dataset) to the validated NeoTCR signature, we observed a strong correlation (r = 0.84), consistent with the previously reported integrated dataset (Figure S4G). In contrast, the corresponding FT-derived gene set (295 genes upregulated in shared versus non-shared TCRs in the FT) showed a more moderate correlation with NeoTCRs (r = 0.65, Figure S4H), suggesting that although tumor-reactive, FT-derived T cells exist in a less exhausted state. The PBMC-derived gene set (117 genes upregulated in shared versus non-shared TCRs in PBMCs) showed a substantially weaker correlation with the NeoTCR signature (r = 0.48, Figure S4I), implying that shared TCRs between PBMC and tumor are less likely to be neoantigen-specific. We further compared these gene sets with another published dataset of neoantigen-reactive peripheral blood lymphocytes (neoPBL). Interestingly, while the correlation for the tumor gene set declined slightly (r = 0.66), the FT gene set showed a marginally stronger alignment (r = 0.68), whereas the PBMC gene set remained poorly correlated (r = 0.37, Figure S4J-L). Together, these findings highlight that the FT gene set is more transcriptionally aligned with known neoantigen-reactive profiles from PBMC, further supporting the notion that FT-resident T cells play a meaningful role in early tumor recognition.

Our findings thus far strongly support the notion that TRMLs in the non-cancerous FT are clonally and functionally related to tumor-specific TRMLs obtained from cancer metastasis in the same patient. FT TRMLs appear less exhausted and maintain a more quiescent, monitoring state compared to tumor TRMLs.

### Co-culturing tumor organoids with FT-derived TRMLs reveals tumor-reactive T cells originating from the FT

We next sought to determine whether TRMLs in the FT are indeed tumor-reactive. We tested TRMLs reactivity *in vitro* by co-culturing non-cancerous FT T cells with either omentum tumor organoids or non-cancerous FT organoids obtained from the same patient. Autologous FT TRMLs were polyclonally pre-expanded from dissociated FT tissue, by co-culture with irradiated PBMCs from 3 healthy donors in combination with exogenous IL-2 and anti-CD28(Figure 5A).^54^ Stimulation of FT TRMLs by tumor organoids invoked a significant induction of IFN-γ response in ELispot assays (178 spots per 10^6^ TRMLs for organoids *vs* 17 spots per 10^6^ cells for FT TRMLs, more than a 10-fold increase for tumor organoids as compared to FT TRMLs, p < 0.0003, Student’s t test, Figure 5B, S5). IFN-γ response by FT TRMLs to autologous non-cancerous FT organoids was undetectable (Figure 5B, S5). To rule out the dependency of tumor organoid- reactive TRMLs on IFN-γ^55^, we compared the expression of the activation marker CD137 by TRMLs upon stimulation with tumor organoids that were either pre-stimulated with IFN-γ or left untreated (Figure S6). From this, we found 14.5% CD137 positive T cells compared to 0.47% positive T cells when cultured with tumor organoids or FT organoids, respectively (p<0.0245, Student’s t test). Together, these data suggest that induced T cell responses were specific for tumor antigens and that FT TRMLs are capable of secondary memory recall upon encountering tumor specific neoantigens expressed by tumor organoids. This result is consistent with the high overlap rate of TCRs between FT and tumor found in the scTCR analysis.

**Figure 5.**
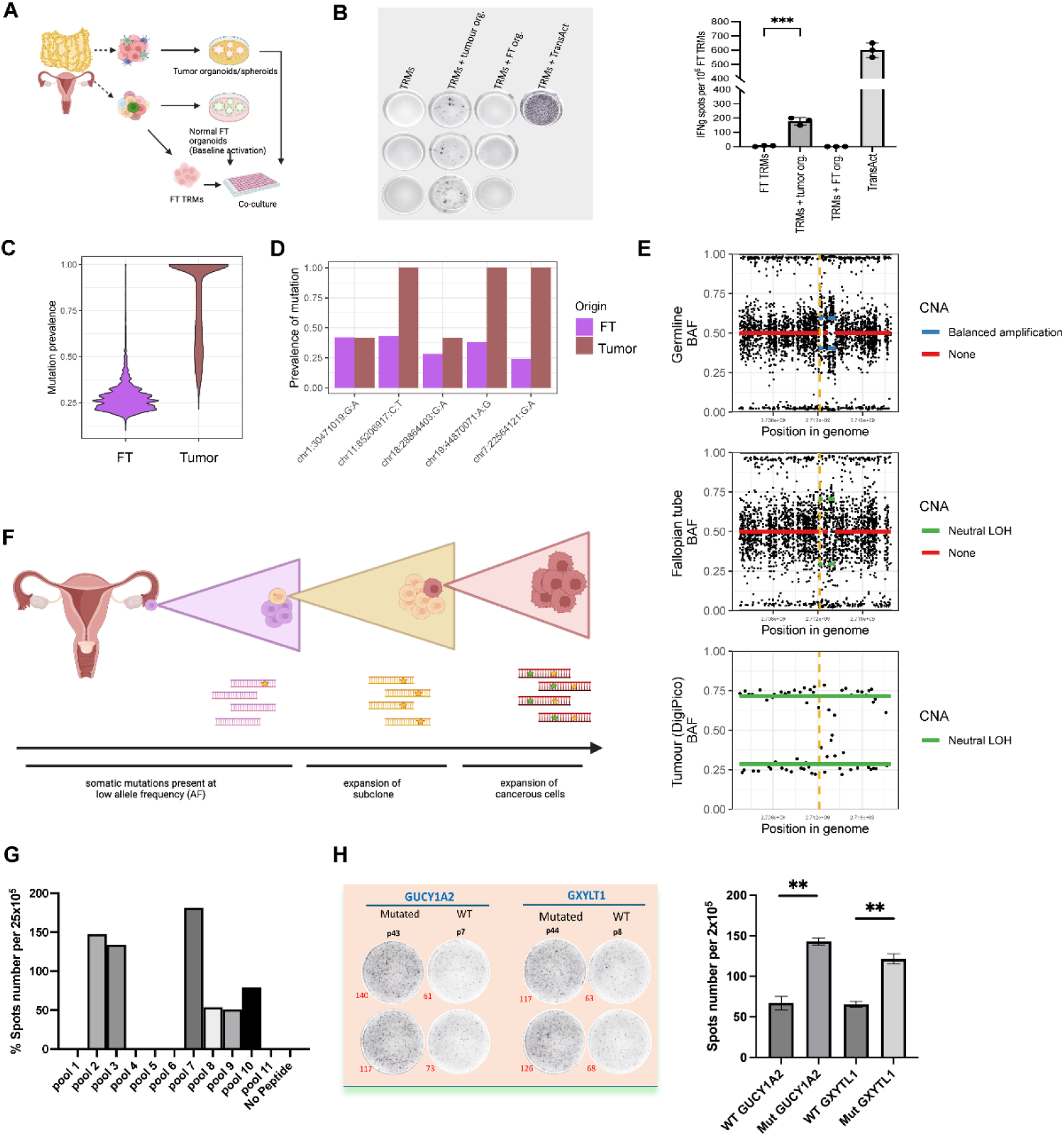
TRMLs from the non-cancerous FT elicit an IFNγ response when co-cultured with autologous metastasis-derived ovarian cancer organoids or tumor mutation-predicted neoantigen peptides. (A) The experimental workflow outlines the establishment of tumor organoids from ovarian cancer resections of omental metastases, which were stimulated with IFNγ for 24 hours before co-culture with FT TRMLs pre- expanded from the same patient. TRMLs were stimulated overnight with tumor or normal FT organoids, and T cell responses were assessed using an IFNγ ELISpot assay. (B) The left panel presents representative well images of the ELISpot plate (see Supplementary Figure S5), while the right panel shows IFNγ ELISpot counts from FT TRMLs of patient 11642 after ex vivo stimulation with tumor organoids or FT organoids. FT TRMLs stimulated with TransAct nanobeads served as a positive assay control. Data are displayed as individual values and the mean of three technical replicates ± SEM. ***p < 0.003 by Student’s t-test, Prism. Data is representative of three experiments with similar results. (C) Prevalence of genome-wide mutations in FT and tumor, computed using OncoPhase. (D) Barplots of the prevalence of the five shared mutations between FT and tumor tissues in corresponding samples. (E) Development of copy number alterations (CNA) at the *PPP2R1A* locus (marked by the yellow dashed line) from germline (top) to FT (middle) to tumor (bottom). CNAs at *PPP2R1A* are known drivers of ovarian cancer. Tumor B allele frequencies (BAFs) are derived from DigiPico data, therefore data displayed here represents de-noise haplotype-averaged BAFs. Germline and FT BAFs were derived from WGS data, therefore each data point here represents a single SNP. (F) Diagrams depicting the progression model of ovarian cancer development. (G) A representative analysis of 27 mutations, organized into eleven pools of 9-mer peptides, selected for immunogenicity testing by ELISpot. The control represents the negative control, and normalized values are plotted (value minus negative control). (H) The left panel shows a deconvolution plot of the pool responses to single peptides, indicating that 2 out of 27 neoantigen candidates (7%) were recognized by pre-existing TRMLs in the FT. The right panel presents a bar plot quantifying the number of spots, with data represented as mean ± SEM. **P < 0.008 by Student’s t-test, Prism.

### Predicted tumor neoantigens elicit FT TRMLs responses

To further characterise this response and to identify putative tumor antigens to which FT TRMLs reacted, we opted to perform detailed potential neoantigen identification using next generation sequencing. Genomic analysis of the tumor showed the typical recognised features of HGSOC including profound genomic instability, *TP53* mutation, *BRCA1* mutation, and a mutation in *NF1*. The genome also harboured known driver copy number alterations (CNAs) found in ovarian cancer, spanning regions including *BRCA1*, *BRCA2*, *KRAS*, *NF1*, *PPP2R1A,* and *TP53* (Figure S7A-C). Mutation calling in the FT organoids revealed low-prevalence somatic mutations compared to those present in the tumor (Figure 5C). There were no somatic mutations in coding regions that were shared between the tumor sample and the FT organoids; however, 5 somatic, non-coding mutations were present in both samples (Figure 5D, see Methods; Mutation overlap). Both the tumor and the FT organoids also showed copy-neutral loss of heterozygosity (LOH) at the *PPP2R1A* locus, which is a known driver of chromosome copy numeric aberration (CNA). In germline PBMCs, this locus showed evidence of triploidy (Figure 5E). We therefore reason that the non-duplicate allele was lost in the FT epithelium prior to tumor formation, resulting in a copy-neutral LOH at that locus (Figure 5E). These shared genomic features suggested that the FT was the tissue-of-origin for the tumor (Figure 5F). Interestingly, the prevalence of three of the shared mutations increased from less than 0.5 in the FT to 1 in the tumor (see Methods; Mutation prevalence) (Figure 5D). This indicated that the sample of cells from the FT may include the cell-of-origin of the tumor. This result further explained the reason for high overlap TCRs of FT and tumor, and it may indicate these TCRs are reactive to the same truncal tumor mutations.

To identify potential expressed tumor neoantigens, we next complemented our analysis with RNA sequencing (RNA-seq) of the tumor. Neoantigen prediction and annotation (see Methods; Neoantigen prediction) identified 41 potential neoantigens spanning 37 mutant epitopes, all of which were expressed in the bulk tumor and predicted to bind MHC-I with high affinity. Twenty seven of these mutations were found to be truncal and were considered as the final set of candidate neoantigens. These 27 predicted neoantigens were divided into pools of 2,5, or 6 peptides (designated as pools 1-11) and were selected for immunogenicity testing (Figure S7D). Six of the pools (45%) were immunogenic, with p5–21, p17–31, p78– 92, and p156–170 eliciting the strongest IFN-γ response (Figure 5G, S7E-F). Three of the 10 peptide pools tested for this patient elicited the strongest autologous T cell responses (Figure 5G). Deconvolution of the pool responses to single peptides showed that 2 of the 27 neoantigen candidate peptides identified by our pipeline (7%) stimulated FT TRM cell responses compared with the matched wild type peptide controls (numbers of spots per 2x10^5^ TRMLs were 143 in mutated GUCY1A2, vs 65 in wt counterpart, and 122 in mutated GXYTL1 vs 65 in its WT counterpart, p<0.007, Student’s t test, Figure 5H).

We next reasoned that for these mutant peptide reactive cells to be biologically relevant, they should possess TCRs that were shared with those obtained by sequencing the tumor organoid reactive T cells. We, therefore, performed ScTCR and ScRNA-seq analysis of FT T cells that were cultured with either tumor organoids or mutant peptides (Figure 6A). We restricted our analysis to CD137+ T cells obtained by FACS sorting as these were presumed reactive T cells. We observed that these tumor-reactive FT T cells exhibited a low number of unique clones while demonstrating high clonal expansion (Figure 6B-C). We identified 14 clones of reactive T cells that were shared between tumor organoid-reactive cells and mutant peptide 43- reactive cells, as well as 15 clones shared between tumor organoid-reactive cells and mutant peptide 44- reactive cells (Figure 6D), which indicates that these reactive T cells are stimulated by similar epitopes. Remarkably, we found that the TCRs of reactive T cells that were co-cultured with tumor organoids, mutant peptide 43 and mutant peptide 44 were shared with 3, 4, and 5 TCRs, respectively from TRMLs in the tumor, and 5, 7 and 7 TCRs from TRMLs in the FT (Figure 6D-F). These findings confirm that T cells in the non-cancerous FT do indeed react to tumor neoantigens.

**Figure 6.**
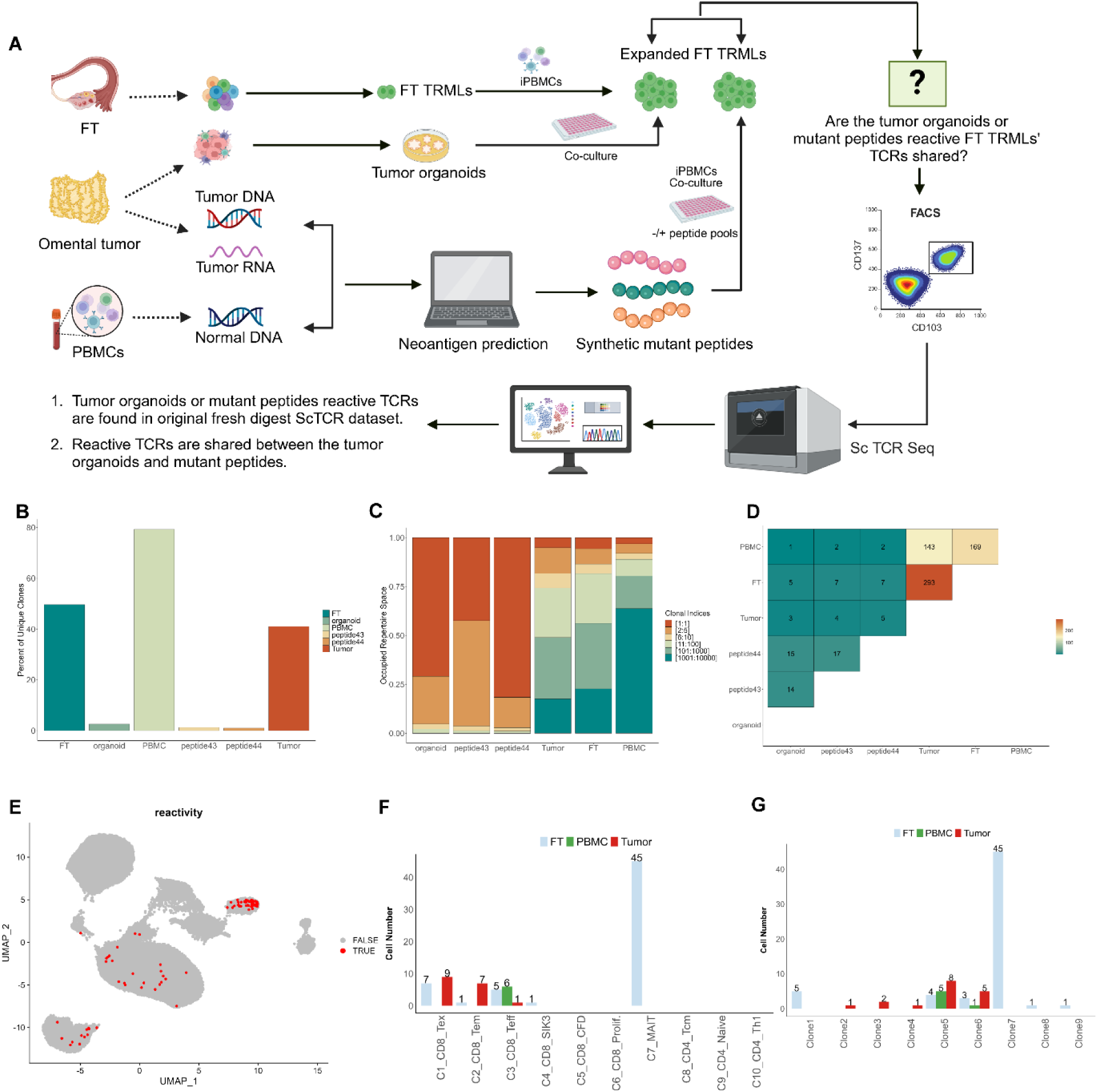
Analysis of T cell clonotypes responsive to predicted tumor neoantigens reveals activation of FT– derived TRMLs (A) Schematic overview of the experimental workflow to identify neoantigen-reactive TCRs. Resected tumor samples were used for tumor organoid culture and for whole-genome and RNA sequencing to identify tumor-specific mutations and predict neoantigens. FT-derived TRMLs were expanded in vitro and co-cultured with tumor organoids or predicted mutant peptides. Activated T cells, identified by CD137 (4-1BB) upregulation, were single-cell sorted and subjected to scTCR-seq. The resulting TCRs were traced back to the original scTCR dataset, revealing 82 cells present in the original FT, PBMC, or tumor samples. Tumor-reactive TCRs were shared between co-cultures with tumor organoids and mutant peptides, indicating recognition of similar antigens. (B) Bar plot showing the percentage of unique clonotypes identified in tumor-reactive FT TRMLs following stimulation with tumor organoids or predicted peptides 43 and 44, compared to the original dataset. (C) Clonal abundance of tumor-reactive FT TRMLs in response to stimulation by tumor organoids and predicted peptides. (D) Clonal overlap plot indicating the number of shared TCR clonotypes across different stimulation conditions and their presence in the original dataset. (E) UMAP visualization showing the distribution of tumor organoid- and peptide 43/44-reactive FT TRMLs (red) within the original dataset. (F) Bar plot displaying the number of tumor-reactive FT TRML cells found across different tissue-derived clusters (FT: blue, PBMC: green, tumor: red) in the original dataset. (G) Bar plot showing the number of tumor-reactive FT TRML clonotypes present across FT, PBMC, and tumor tissues in the original dataset, supporting their immunological relevance.

Cells with shared TCRs were sparsely distributed across clusters C1_CD8-Tex, C2_CD8-Tem, C3_CD8- Teff, C6-CD8-Prolif., C7-MAIT (Figure 6E-F), indicating that under-expanded TCRs in patients may also be tumor-reactive and can be successfully expanded *in vitro*. We further traced tumor-reactive FT TCRs in our original dataset. There are 2 TCRs that can be found in FT, tumor, and PBMC (Figure 6G). Interestingly, there are 3 clones unique to the original tumor dataset (Figure 6G), but we used the FT T cells for the assay, which implies these TCRs were also shared with FT. Therefore, 5 TCRs in total are shared with FT and tumor in this assay, which further supports our hypothesis that T cells from FT and tumor are recognising the same tumor antigens.

## DISCUSSION

This work provides a comprehensive single-cell RNA sequencing (scRNA-seq) and paired single-cell T cell receptor sequencing (scTCR-seq) analysis of TRMLs from the FT, omental tumor metastases, and peripheral blood in HGSOC patients. We found that FT TRMLs exhibit substantial TCR overlap with TRM-like TILs, with 373 out of 2,029 TCR clonotypes shared between the two sites. This overlap was significantly higher than that observed between peripheral blood and tumors, suggesting that a subset of FT TRMLs serves as a precursor immune surveillance population capable of recognizing tumor-derived neoantigens before metastatic dissemination. The functional significance of these shared TCRs was supported by *in vitro* experiments demonstrating that FT TRMLs exhibit tumor reactivity, as evidenced by their response to tumor organoids and tumor-derived peptides. These findings support the notion that TRMLs in the FT are actively engaged in immune surveillance against early tumor lesions, a process that could be leveraged for cancer prevention and immunotherapy development. The high degree of TCR overlap between FT TRMLs and tumor TRMLs supports a model in which tumor-reactive T cells originate in the FT before tumor dissemination. The fact that shared clonotypes were predominantly enriched in exhausted and proliferative exhausted CD8+ subsets suggests that these cells have undergone antigen-driven expansion, providing evidence for early immune engagement with tumor antigens. This phenomenon aligns with models of TRM-mediated immune equilibrium in epithelial malignancies, such as cervical cancer, where TRMs restrict HPV-driven transformation until immune escape occurs^16^, and early-stage melanoma, where TRMs limit tumor progression by responding to neoantigens at premalignant stages.^56^ Our findings suggest a similar mechanism in HGSOC, where the FT serves as an immune surveillance site, keeping preneoplastic lesions in check before eventual immune evasion and tumor progression. The increased expression of immune checkpoints (*CTLA4*, *LAG3*, *PDCD1*, *TIGIT*) in tumor TRMLs compared to FT TRMLs suggests that chronic antigen stimulation and the tumor microenvironment drive exhaustion in these cells. The presence of exhaustion markers in tumor-infiltrating TRMLs, but not in their FT counterparts, suggests that TRMLs in the FT are functionally more potent and may represent a better source of tumor-reactive T cells for immunotherapy. This raises the possibility that T cells isolated from FT tissue could be used in adoptive T cell therapy or engineered to resist exhaustion, enhancing their anti-tumor potential.

Based on our findings, we propose a model to explain the high degree of TCR overlap between TRMLs in the FT and the tumor, as well as the tumor-reactive capacity of FT-derived TRMLs. In the early precancerous stage, when the FT epithelium first encounters tumor-associated antigens, distant dendritic cells (DCs) are recruited and migrate to the site of origin. These DCs process and present tumor-associated antigens (TAA) to naïve T cells in the pelvic lymph nodes (LN), thereby priming an immune response against tumor cells that express both normal and mutated peptides. This interaction leads to the formation of memory T cells that remain in the LN until subsequent exposure to the same antigen. Upon re-stimulation with their cognate antigen, these memory T cells migrate to sites of antigen-expressing lesions, regardless of their location, where they undergo proliferation and mediate cytotoxic responses against tumor cells. These findings reinforce the hypothesis that HGSOC originates in the FT and provide a new perspective on the role of TRMLs in regulating immune surveillance and maintaining immune equilibrium at the earliest stages of ovarian cancer development. A major implication of this model is the potential for developing vaccines for ovarian cancer prevention. Given that FT TRMLs exhibit tumor-reactive properties at an early stage of disease progression, they could serve as an ideal target for immunization strategies aimed at boosting pre-malignant immune responses before full tumor development. The high TCR overlap between FT and tumor sites further suggests that tumor-reactive immune responses originate early in the disease course, highlighting the potential for preemptive immunotherapeutic interventions.

By specifically targeting TRMLs for sequencing, we successfully identified and annotated a novel TRM C4_CD8-SIK3 cluster. We had previously reported that a close family member SIK2 was a key regulator of ovarian cancer omental metastasis and a target for therapy^57–60^ and found that it was activated by increased cytoplasmic calcium concentration in cancer cells.^58^ A rise in calcium concentration is a key feature following T cell engagement.^61^ We further validated the SIK3 and CD8 co-expression by immunofluorescence staining. To our knowledge, this is the first evidence of SIK3 expression by CD8 TRMLs in tumor and FT tissues. The enrichment of genes involved in cell differentiation suggests that C4_CD8-SIK3 cells are either in a state of differentiation or play a crucial role in maintaining a specialized T cell phenotype. The enrichment of histone modification pathways in *SIK3*-expressing TRMLs suggests that these cells may possess a unique degree of epigenetic plasticity that enables them to function in the tumor microenvironment despite chronic antigen stimulation. Given that epigenetic remodelling is a key hallmark of T cell exhaustion, the identification of SIK3-high TRMLs suggests that this population may represent a transition state between functional and exhausted T cells. Targeting SIK3-dependent pathways could provide a new strategy for reprogramming dysfunctional T cells to restore their anti-tumor capacity in ovarian cancer. Further functional studies are warranted to explore the specific roles of these pathways in the context of T cell-mediated immunity against STICs, where T cell function is a critical determinant of preventive and therapeutic intervention.

Our findings provide key insights into the immunobiology of ovarian cancer and highlight potential strategies for improving immunotherapy, which has thus far shown limited efficacy in this malignancy. The failure of current immunotherapeutic approaches in ovarian cancer can be attributed to several factors, including the presence of dominant immunosuppressive signals that inhibit T cell reactivation^62^, metabolic constraints imposed by the tumor microenvironment and the peritoneal cavity^49^, and a substantial accumulation of immunosuppressive myeloid cells^63,64^, ovarian cancer’s heterogeneity and tumor mutates to downregulate the immunogenic antigen and thus evades immune system surveillance and continues to proliferate.^65–67^ These challenges emphasize the urgent need for novel strategies to counteract immune suppression and enhance therapeutic responses in ovarian cancer. One of the most striking findings in this study is the high degree of TCR overlap between TRMLs in the FT and TILs. We found that these shared clonotypes were predominantly enriched in the C1_CD8-Tex cluster in the FT and the C1_CD8-Tex, C2_CD8-Tem, and C6_CD8-Prolif. clusters within tumors. This supports recent studies suggesting that TRM-like precursor cells may differentiate into terminally exhausted cells as they encounter chronic antigen stimulation in the tumor microenvironment.^68^ Furthermore, our findings reveal that CD8+ TRM-like TILs in the tumor and circulating PBMCs represent highly distinct compartments in terms of phenotype and function. The low degree of TCR overlap between tumors and PBMCs, with shared clonotypes being primarily restricted to the Teff cluster in PBMCs, suggests that most circulating T cells are bystanders and do not contribute significantly to tumor immune surveillance. Crucially, we demonstrate that TRMLs from the FT retain the capacity to recognize tumor antigens, suggesting that these cells may represent an untapped resource for immunotherapeutic intervention. Given their less exhausted phenotype and tumor-reactive capacity, it is plausible that FT-derived TRMLs could serve as the basis for personalized adoptive T cell therapies. Instead of relying on highly dysfunctional TILs, TCR-engineered T cells derived from FT TRMLs could be expanded and reintroduced into patients, potentially overcoming the challenge of antigenic heterogeneity across metastatic sites. These findings open new possibilities for harnessing the immunological memory of FT TRMLs in a personalized manner, offering a promising avenue for improving immune-based therapies in ovarian cancer.

In conclusion, this study represents the first comprehensive characterization of TRMLs in the FT and their role in ovarian cancer progression. Our findings establish the FT as an immune surveillance site, revealing that FT TRMLs recognize tumor neoantigens, exhibit clonal expansion, and are less exhausted than tumor-infiltrating T cells. The discovery of a novel SIK3-high TRML subset further suggests that epigenetic and metabolic plasticity may play a role in maintaining immune equilibrium during early tumorigenesis. These findings provide an important foundation for developing next-generation immunotherapies and cancer prevention strategies that leverage the natural tumor-reactive capacity of TRMLs in ovarian cancer.

## ACKNOWLEDGMENTS

We thank the WIMM Flow Cytometry Facility, Single Cell Facility, and CCB for their help in this study. This work was supported by Ovarian Cancer Action. T.T is funded by Kanzawa Medical Research Foundation, The Watanabe Foundation, The Yasuda Medical Foundation, and JSPS Overseas Research Fellowships.

## AUTHOR CONTRIBUTIONS

Conceptualization, Project Administration, and Supervision, A.A.A.; Funding Acquisition, A.A.A.; Investigation, L.W., B.H., N.H., J.N., A.A., M.S., L.L.W., L.R., A.A.E., A.A.D., T.T., M.A., H.S.M., J.Y., C.Y., N.Z., and A.A.A.; Data Curation, A.A.A., C.Y., N.Z., and L.W.; Methodology and Formal Analysis and Visualization, A.A.A., L.W., and N.Z.; Writing Original Draft, A.A.A., L.W., N.Z., and B.H.; Writing – Review & Editing, A.A.A., L.W., N.Z., and C.Y.; Resources (Clinical Samples), H.S.M., and J.Y.

## STAR★METHODS

### KEY RESOURCES TABLE

| REAGENT OR RESOURCE | SOURCE | IDENTIFIER |
| --- | --- | --- |
| Antibodies |  |  |
| Anti-SIK3, polyclonal, antibody produced in rabbit | Abcam | Cat#ab110987 |
| Anti-CD8, Monoclonal, Clone # 1033123, produced in mouse | R&D | Cat#MAB3803-SP |
| Anti-CD3, Clone OKT3, BV711 conjugated antibody | BioLegend | Cat#317328<br>RRID:AB11219592 |
| Anti-CD103, Ber-ACT8, PerCP/Cyanine5.5 conjugated antibody | BioLegend | Cat#350226<br>RRID:AB_2716191 |
| Anti-CD4, Clone A161A1, PE-Cy7 conjugated antibody | BioLegend | Cat#357410<br>RRID:AB_2565662 |
| Anti-CD8a, Clone RPA-T8, APC conjugated | BioLegend | Cat#301014 RRID:<br>AB_2562054 |
| Anti-CD137, Clone 4B4-1, PE conjugated | BioLegend | Cat#309804<br>RRID:AB_10933428 |
| DAPI | Miltenyi Biotec Ltd | Cat#130-111-570 |
| Anti-CD56, Clone HCD56, PE conjugated | BioLegend | Cat#318305 RRID:AB_604101 |
| Anti-Epcam, Clone 9C4, FITC conjugated | BioLegend | Cat#324210 RRID:AB_756083 |
| Anti-CD137 (4-1BB), Clone 4B4-1, APC/Cyanine7 conjugated antibody | BioLegend | Cat#309830 RRID<br>AB_2629644 |
| Mouse anti-human CD28 | eBioscience | Cat#16-0289-81,<br>RRID:AB_468926 |
| Anti-human PD1 / nivolumab | Merus / Selleckchem | Cat#A2002 RRID:AB_2810223 |
| CD3 Antibody, anti-human, (OKT3) | Miltenyi Biotec Ltd | Cat#130-093-387<br>RRID:AB_1036144 |
| Chemicals, peptides, and recombinant proteins |  |  |
| IMDM, GlutaMAX Supplement | Life Technologies Ltd | Cat#31980048 |
| Penicillin-Streptomycin (10,000 U/mL)-100 mL | Life Technologies Ltd | Cat#15140122 |
| Amphotericin B | Life Technologies Ltd | Cat#15290026 |
| Lonza Gentamicin 50 mg/ml | Scientific Laboratory<br>Supplies Limited | Cat#LZ17-518Z |
| HUMAN SERUM TYPE AB (MALE) FROM CLOTTED | Merck Life Science UK<br>Limited | Cat#H5667-100ML |
| Human IFN-gamma ELISpotPRO kit (ALP) | Mabtech AB | Cat#3420-2AST-2 |
| TransAct human | Miltenyi Biotec Ltd | Cat#130-111-160 |
| FcR Blocking Reagent, human | Miltenyi Biotec Ltd | Cat#130-059-901 |
| recombinant human IFN gamma | Peprotech | Cat#300-02 (100ug) |
| DPBS | ThermoFisher | Cat#14190144 |
| Trypsin-EDTA (0.5%), no phenol red-100 mL | ThermoFisher | Cat#15400054 |
| Advanced DMEM/F-12 | ThermoFisher | Cat#12634010 |
| EGF | ThermoFisher | Cat#PHG 0311 (100µg) |
| human Noggin | Peprotech | Cat#120-10C (5µg) |
| human Fibroblast Growth Factor-10 (FGF-10) | Peprotech | Cat#AF-100-26 (5µg) |
| N2 supplement 100X | ThermoFisher | Cat#17502048 (5ml) |
| B27 supplemented50X | ThermoFisher | Cat#17504044 (10ml) |
| Nicotinamide | Sigma-Aldrich | Cat#N0636-100G (100g) |
| N-acetyl L-Cysteine | Sigma-Aldrich | Cat#A9165-5G |
| A83-01 | Tocris | Cat#2939/10 (10mg) |
| Forskolin | Tocris | Cat#1099/10 |
| Y27G32 | Tocris | Cat#72302 (1mg) |
| DNase I | Roche | Cat#4716728001 |
| Collagenase Type 1 | Invitrogen | Cat#17018029 |
| matrigels matrix phenol red free mouse ehs tumor 10ml | Corning | Cat#356237 |
| DMSO | Sigma | Cat#sc-358801 |
| Fetal Bovine Serum, qualified, heat inactivated, Brazil | ThermoFisher | Cat#10500-064 |
| Tumor dissociation kit, human (contains Enzymes H, R, and A) | Miltenyi Biotec Ltd | Cat#130-095-929 |
| Critical commercial assays |  |  |
| SpectraMax® Quant™ AccuBlue™ Pico dsDNA Assay Kit | SpectraMax® | Cat#R8355 |
| REPLI-g Advanced DNA Single Cell Kit | Qiagen | Cat#150363;24 samples |
| SMARTer Stranded Total RNA-Seq kit v2 - Pico Input | Takara | Cat#634412 |
| GenElute Single Cell RNA Purification Kit | Sigma-Aldrich | Cat#RNB300 |
| Chromium Next GEM Single Cell 5' Kit v2, 4 rxns | 10x Genomics | 1000265 |
| Library Construction Kit, 16 rxns | 10x Genomics | 1000190 |
| Dual Index Kit TT Set A, 96 rxn | 10x Genomics | 1000215 |
| Chromium Single Cell Human TCR Amplification Kit, 16 rxns | 10x Genomics | 1000252 |
| Chromium™ Next GEM Chip K Single Cell Kit, 16 rxns | 10x Genomics | 1000287 |
| Software and algorithms |  |  |
| R 4.2.3 | R Core Team | https://www.r-project.org/ |
| Cellranger-7.0.0 | 10x Genomics | https://www.10xgenomics.com/ |
| Seurat v4.0 (R package) | <sup>69</sup> | https://satijalab.org/seurat/ |
| scRepertoire 2 | <sup>70</sup> | https://www.borch.dev/uploads/sc repertoire/ |
| Monocle 3 | <sup>71-73</sup> | https://cole-trapnell-lab.github.io/monocle3/ |
| Harmony (R package) | <sup>74</sup> | https://github.com/immunogenomics/harmony |

### RESOURCE AVAILABILITY

#### Lead contact

Further information and requests for resources should be directed to and will be fulfilled by the Lead Contact, Ahmed Ahmed. This study did not generate new unique reagents.

#### Materials availability

All reagents are commercially available.

#### Data and code availability

All software used in this study is freely available. Any additional information including custom code required to analyze the data in this study will be available upon request.

## EXPERIMENTAL MODEL AND STUDY PARTICIPANT DETAILS

### Human Fallopian Tube and Tumor Samples

The cases in this study were recruited under the Gynecological Oncology Targeted Therapy Study 01 (GO- Target-01). All participants involved in this study were appropriately informed and consented. Fallopian tube (FT) biopsies for single-cell RNA-seq were collected from the distal end of fallopian tubes. In total, we collected FT and omentum samples from four patients with HGSOC.

## METHOD DETAILS

### Ovarian cancer digestion and 3D culture protocol

Tumor tissue samples were set on a sterile petri dish on crushed ice. Identifiable necrotic tissue and fat tissue were removed as much as possible, and their size and weight were measured. Ovarian tumor tissues were cut into small pieces of around 2-4 mm and incubated in prewarmed RPMI 1640, 1% Penicillin/Streptomycin (Gibco), transferred into gentleMACS C Tube containing the enzyme mix. Tumor pieces were dissociated at 37 °C using the Human Tumor Dissociation kit (Miltenyi Biotec) on a gentleMACS Octo Dissociator. After dissociation, single-cell suspensions were filtered and washed with ammonium-chloride-potassium (ACK) lysing buffer. Cell counts and viability will be assessed after staining with trypan blue. 3D ovarian tumor culture was carried out as previously described^75,76^ with some modification. Organoid culture: 1.0 × 10^4^ cells were then suspended in Matrigel (Corning), 30 μL drops of matrix cell suspension is allowed to solidify on a pre-warmed 24-well plate at 37 °C for 20 min. After polymerization of the Matrigel, organoid medium cocktail was added. Advanced DMEM/F12 (Thermo Fisher Scientific) was supplemented with 10 mM HEPES (Thermo Fisher Scientific), 1 × GlutaMAX-I (Thermo Fisher Scientific), 1X B27 supplement (Thermo Fisher Scientific), 1X N2 supplement, 1 mM N-acetylcystein (Sigma-Aldrich), 1mM Nicotinamide, 100 ng/mL recombinant human EGF 10 ng ml-1 (R&D Systems), 50 ng/mL recombinant human FGF-2 (PeproTech), 1 µg/mL humanR-spondin (R&D Systems), 100 ng/mL Noggin (PeproTech), 500 nM A-83-01 (Tocris Bioscience), 200 U/mL penicillin/streptomycin (Thermo Fisher Scientific), and 10 µM Y-27632 (Tocris Bioscience). The medium was changed every 3–4 days, and the organoids were passaged at a 1:3 dilution every 1–4 weeks.

### FT digestion and organoids culture protocol

FT tissue samples were set on a sterile petri dish on crushed ice, and were washed 3x times with cold DPBS and discard supernatant (ensure removal of excess Red Blood Cells). FT tissues were cut into small pieces and place tissue in pre-warmed “Digestion Medium”: DMEM R0 (RPMI 1640 + 1% P/S), 0.5 mg/ml DNase I and 100 Units/ml Collagenase Type I). Enzymes should be kept in working aliquots at -20°C and added to the medium just before use. FT tissues were cut again into very small pieces, and collected with a 50 ml pipette into a 15 ml Falcon tube. Digestion medium was added for a total volume of 5ml per C Tube. The tissue pieces were transferred into the gentleMACS C Tube containing the enzyme mix. The C Tube was closed and attached upside down onto the sleeve of the gentleMACS Dissociator. The 37C_h_TDK_1 gentleMACSTM Program was selected. After dissociation, single-cell suspensions were filtered and washed with ammonium-chloride-potassium (ACK) lysing buffer. Cell counts and viability will be assessed after staining with trypan blue. 0.1M of FT tissue digest were seeded in 30 μl of Matrigel (Corning) in pre-warmed 12-well plates. The Matrigel was solidified for 20 min at 37 °C and overlaid with 1 ml pre-warmed expansion medium (DMEM/F12 with 500 ng ml-1 of human recombinant R-Spondin-1, supplemented with 12 mM HEPES, 1% GlutaMAX, 2% B27, 1% N2, 10 ng ml−1, human EGF (Invitrogen), 100 ng ml−1 human noggin, 100 ng ml−1 human FGF10 (all from Peprotech), 1 mM nicotinamide, 9 μM ROCK inhibitor (Y- 27632, both from Sigma), 5 μM TGF-β TGF-β type I Inhibitor (A83-01, Tocris) and 10 µM Forskolin (R&D systems). Cultures were kept at 37 °C, 5% CO2 in a humidified incubator. A tissue fragment of ∼5 cm yields ∼8 × 10^6^ cells, within 7 days post isolation, this yields almost 10^8^ cells within 4 weeks of isolation.

### Expansion of FT TRMLs

FT T cells were generated from single cell suspensions of digested primary FT tissues and expanded as described previously ^77^ with slight modifications. Briefly, FT T cells were cocultured with irradiated (70 Gy) healthy donors’ PBMC in a ratio of 1:200 in IMDM complete medium (REP) (10% human AB serum, 3000 IU/mL interleukin 2 (Proleukin, Novartis, Basel, Switzerland), 10 μg/ml gentamicin, 100 U/ml penicillin and 100 μg/ml streptomycin, 1.25 μg/ml amphotericin B (Fungizone), supplemented with 30µg/ml OKT3 (miltenyi biotec). On day 5 of coculture, 3⁄4 of the medium was replaced with fresh REP medium. Starting at day 7, cells were counted daily, and cell density was adjusted to a concentration of 1.0×10^6^ cells/mL REP medium. Expansion will be ended at day 14 and expanded cells will be frozen or directly used for further experiments.

### FACS sorting

Following tissue enzymatic dissociation of tissues, single-cell suspensions were subjected to antibody staining with anti-EPCAM, anti-CD56, anti-CD3 and anti-CD103. Cells were purified by fluorescence-activated cell sorting (FACS) on SONY MA400 cell sorter equipped with four lasers (405 nm, 488 nm, 561 nm and 638nm). A 100-μm nozzle running at 70 psi and 90 kHz was used as the setup for each sort session.

FlowJo (v.10) was used to collect and analyze the flow cytometry data. Before gating on fluorescence, single cells were gated using forward scatter (FSC-A) and side scatter (SSC-A) (for intact cells) and SSC-W/SSC-H and FSC-W/FSC-H (to ensure that only singlets were sorted). FACS gates were drawn to include only live single cells based on DAPI (Thermo Fisher Scientific). Further gates were drawn to arrive at CD3+CD103+EpCAM− (for CD103+selected samples) or CD3-EpCAM+ cells (for EPCAM+ selected samples). Boundaries between positive and negative cell fractions were drawn based on single-colour stains. An example gating strategy is shown in Supplementary Methods (Figure S1).

### Generation and detection of patient neoantigen-specific FT TRMs cells ELISPOT assays

Prior to co-culture with autologous FT TRMs cells, tumor or FT organoids were pre-stimulated with IFNγ to enhance antigen presentation. IFNγ exposure also led to the induction of PD-L1, a negative regulator of T cell activation^78^, and to counteract any inhibitory effect of PD-L1 during T cell activation, blocking antibodies to PD-1 were added.^79^ For T cell response and specificity to tumor organoids, FT TRMLs were cocultured overnight with autologous tumor organoids or FT non-cancerous organoids in IFNγ ELISPOT plate in IMDM medium supplemented with penicillin–streptomycin and 10% AB human serum (complete IMDM) as previously described.^79^ For neoantigen specific T cell response and deconvolution of T cell responses, 27 peptides were selected and screened ex vivo to evaluate their immunogenicity against FT TRMs. Pools of these peptides were incubated with FT TRMs at 0.25 × 10^6^ cells per well with individual (2μg/ml) or pooled peptides (each at 2μg/ml) in the presence of IL-2 (3000 IU/ml; Proleukin), cocultured with allogeneic irradiated PBMCs for 14 days before performing IFN-γ ELISPOT assays using no peptide and TransAct stimulation as negative and positive controls, respectively. After this, viable cells were restimulated for 18- h with or without peptides in an ELISPOT plate and analysed for interferon-γ (IFN-γ) production in response to the individual peptides comprising the pool. The averages of responses to DMSO for each time point were subtracted from experimental wells for background correction. Upon detection of a positive ELISPOT response (defined as at least 55 spot-forming units (SFU) per 10^6^ T cells or a ≥3-fold increase over baseline), the peptide pools were allocated into sub-pools to deconvolute the peptide pool to determine the immunogenic peptide(s) per pool.

### Immunofluorescence and confocal microscopy

CD8 TILs isolated from patients with HGSOC were stained with anti-CD8 antibody and FACS-sorted, then subjected to immunofluorescence and evaluated with confocal microscopy. Cells were fixed using 4% paraformaldehyde/PBS for 10 min, permeabilized in 0.1% TX-100/PBS for 8 min, and incubated in a blocking solution (10% BSA in PBS) for 2 hr at RT. Cells were then incubated with the following primary antibodies at 1/200: Anti-SIK3 antibody (rabbit) ab110987, Abcam and Anti-CD8 (mouse) MAB3803-SP, R&D, overnight at 4℃, washed in PBS (3 times) and incubated in Alexa fluor conjugated-secondary antibodies for 2 hr at RT. The cells were then washed with PBS, mounted in ProLong™ Gold Antifade Mounting (P36930, ThermoFisher Scientific), and imaged using a Zeiss confocal microscope. All images were acquired at 1024-pixel resolution with either oil objective or similar settings of the microscope.

### Single cell RNA-seq and VDJ profiling library and sequencing

TRM-like and circulating T cells were sorted from 4 different HGSOCs for scRNA-seq analysis. Single-cell RNA-sequencing was performed using the 10X Genomics Chromium. A single-cell suspension derived from dissociated tissue was sorted. The cell suspension was loaded onto the 10X Genomics Chromium Single Cell Controller at a concentration of 700∼1000 cells per microliter in order to encapsulate between 5000 and 12,500 cells per sample. Briefly, the single cells, reagents, and 10x Genomics gel beads were encapsulated into individual nanoliter-sized Gelbeads in Emulsion (GEMs) and then reverse transcription of poly-adenylated mRNA was performed inside each droplet. The cDNA and VDJ-enriched libraries were completed in a single bulk reaction using the 10X Genomics Chromium Next GEM Single Cell v2 and V(D)J Reagent Kits. 25,000 or 5,000 sequencing reads per cell for scRNA-seq or VDJ libraries, respectively, were generated on the NovaSeq X Plus instrument. Demultiplexing, barcode processing, alignment, and gene counting were performed using the 10X Genomics CellRanger v7.0 software.

### Single-cell RNA-seq data processing, filtering, batch effect correction, and clustering

Raw sequencing reads from scRNA-seq were processed using Cell Ranger (v7.0, 10X Genomics). Briefly, the base call (BCL) files generated by Illumina sequencers were demultiplexed into fastq files based on the sequences of the sample index and aligned against GRCh38 human transcriptome using STAR.^80^ Cell barcodes and UMIs associated with the aligned reads were subjected to correction and filtering. Filtered gene-barcodes matrices containing only barcodes with UMI counts passing the threshold for cell detection were imported to Seurat v4.2.1^69^ for downstream analysis. We first removed ambient RNA by the default setting of the R library package SoupX_1.6.2^81^, and then removed the doublets by the default setting of the R library package DoubletFinder_2.0.3^82^. Then keep cell barcodes with more than 300 and lower than 6000 genes expressed, lower than 15% UMIs originated from mitochondrial genes, nCount_RNA more than 1000 and lower than the quantile (nCount_RNA,0.97), and lower than 3% hemoglobin gene. This process resulted in 51,855 cells being left. For each sample, standard library size and log-normalization were performed on raw UMI counts using NormalizeData(), and the top 3000 most variable genes were identified by the ‘‘vst’’ method in FindVariableFeatures(). In each study, individual data were further integrated to remove batch effects and integrate data using the RunHarmony function in the harmony library (harmony_1.2.0). Clustering was performed using the FindClusters() function at resolution 0.2 for integrated data (10 clusters). Uniform manifold approximation and projection (UMAP) was used to visualize single-cell gene expression profile and clustering, using the RunUMAP() function in Seurat with default settings.

### Differential gene expression analysis and cluster annotation

Differential expression analysis was performed using the FindAllMarkers() function in Seurat with logfc.threshold=0.25, min.pct=0.1, and test.use=’’wilcox’’. Cells within each cluster were compared against all other cells. Genes with Bonferroni-corrected p-value <0.05 and an average log-fold change > 0.25 were considered differentially expressed. Results from differential gene expression analysis are provided in Table S1. Clusters were annotated by comparing differential genes with markers associated with T cell exhaustion/dysfunction (*PDCD1*, *LAG3*, *TIGI*T, *CTLA4*, *GZMK*, *HLA_DRB1*), effector memory (*CD7*, *XCL1*, *AREG*, *HSPA1B*)), effector (*GNLY*, *FGFBP2*, *GZMH*, *IFIT2*), central memory (*PRKCA*, *FOXP1*), MAIT(*KLRB1*, *IL18R1*, *CCR6*), proliferation (*STMN1*, *MKI67*, *DNMT1*), and Th1 (*TNFRSF4*, *ICAM2*, *CCR7*).

### Visualization of marker genes on singe cell RNA-seq clusters and groups

Marker genes were visualized on UMAP projections or violin plots using log-normalized counts. For the bubble plot of marker genes, the average expression of each gene was calculated for each cluster/group and then normalized by mean and standard deviation.

### scTCR-seq data processing

The TCR sequence data from 10X Genomics were processed using Cell Ranger software (v.7.0) with the manufacturer-supplied human VDJ reference genome. For each sample, the output file filtered_contig_ annotations.csv, containing TCR α- and β-chain CDR3 nucleotide sequences, was used for downstream analysis. Only those assembled chains that were productive, highly confident, full length, with a valid cell barcode and an unambiguous chain type (for example, alpha) assignment were retained. If a cell had two or more qualified chains of the same type, only that chain with the highest UMI count was qualified and retained. For each patient, cells with an identical α/β-chain pair were considered as having originated from the same clonotype and were therefore identified as clonal cells. scRepertoire_2.0.4 package was used to further analyze scTCR-seq data. Shared clones were identified by the ‘‘strict’’ method in clonalOverlap(). Gene set variation analysis (GSVA) was used to calculate the scGSEA score and the correlation between our geneset and the published reference geneset, which were provided in Tables S2.

### Differential expression and Gene Ontology enrichment analysis

The significantly overexpressed marker genes for clusters were identified using Seurat’s FindAllMarkers() function. Genes with adjusted P value < 0.05 by the Wilcoxon rank-sum test were defined as cluster-specific signature genes. For two different clusters, we used the Wilcoxon test to evaluate the significance of each gene, with multiple hypothesis corrections using the BH procedure. Genes with adjusted P value <0.05 were considered as differentially expressed genes that were further used for GO enrichment analysis with the clusterProfiler package (clusterProfiler_4.6.0). GO terms with adjusted P values <0.05, using the BH procedure, were considered significant.

### Building single-cell trajectory

We constructed cell trajectories with Monocle 3 using the top 50 expanded shared TCRs between FT and tumor scRNA-seq data^83^. We normalized data again by Seurat, and the top 2000 most variable genes were identified by the ‘‘vst’’ method in FindVariableFeatures. Data were further integrated to remove batch effects and integrate data using the RunHarmony function using the R package Harmony. Clustering was performed using the FindClusters() function at resolution 0.4 for integrated data. The library SeuratWrappers was used to convert Seurat data to Monocle3 CDS object by function as.cell_data_set(). We chose the cell derived from the Tem cluster as the ‘root’ of the trajectory. Monocle 3 ordered each cell along a learned trajectory according to its transcriptional progress. The default parameters in Monocle 3 were used for the analysis. The exhaustion score was calculated by averaging the expression levels of a predefined set of exhaustion-related genes (*CXCL13*, *CTLA4*, *LAG3*, *HAVCR2*, *TIGIT*, *PDCD1*, *TOX*, *STAT1*) for each cell. Only genes present in the dataset were included, and the score was computed across the selected sample cells.

### Whole-genome sequencing (WGS)

For FT organoids and PBMC samples, the Genomic DNA (gDNA) was extracted with the DNeasy Blood & Tissue Kits (Qiagen), and quantified by Qubit. The WGS libraries were prepared by Novogenes company and sequenced pair-ended on Illumina Novaseq 6000. For the tumor sample, tumor cells were sorted by Epcam+ by FACS, and the DNA was extracted by REPLI-g Advanced DNA Single Cell Kit and quantified by SpectraMax® Quant™ AccuBlue™ Pico dsDNA Assay Kit Cat. We used our in-house method DigiPico^84^ to construct the library and sequenced pair-ended on Illumina Novaseq 6000.

#### WGS data QC & variant-calling

Trimming/alignment QC: Raw FASTQ files were merged across Illumina sequencing lanes and then trimmed using TrimGalore! (version 0.6.7). Trimmed reads were then aligned using bwa-mem2 (version 2.2.1)^85^ and PCR duplicates were removed from the merged BAM file using Sambamba (version 1.0).^86^ Deduplicated BAM sequence calls were updated using base quality score recalibration (BQSR) from GATK (version 4.2).^87^

Variant calling: Candidate variants were called in either the bulk PBMCs or FT organoid using Mutect2 from GATK (version 4.2)^87^ and Strelka (version 2.9.10).^88^ Only consensus mutations called in both were taken forward as the final set of candidate variants. Variants in tumor data were called from the merged DigiPico BAM file against the hg19 reference genome using platypus (version 0.8.1.2), and candidate variants also present in the germline PBMCs were removed from further analysis.^89^ MUTLX was used to determine a final set of mutations as described in.^84^ Final called mutations were phased and passed onto OncoPhase (version 0.1) to determine the mutation prevalence within the tumor sample.^58,90^

Estimating Copy Number Alterations (CNAs): CNAs were estimated using ASCAT (version 2.5.2)^91^, and mutational prevalence was estimated using OncoPhase (version 0.1).^90^ Tumor data CNAs were estimated as described in Nulsen, J. et al (2023).^92^

### Bulk RNA-sequencing

Tumor cells around 5,000-20,000 cells were sorted by Epcam+ by FACS. The RNA was extracted by GenElute Single Cell RNA Purification Kit (Sigma-Aldrich) , and quantified by Agilent RNA 6000 Pico Kit. The SMARTer Stranded Total RNA-Seq kit v2 - Pico Input (Takara) was used to prepare sequencing libraries which were indexed, enriched by 15 cycles of amplification, assessed using the Agilent TapeStation, and then quantified by Qubit. The library was sequenced using 150 bp pair-end reads on the Illumina Novaseq 6000 platform.

Trimming, alignment & quantification: Raw FASTQ files were first merged across Illumina sequencing lanes and replicates, then trimmed using TrimGalore! (version 0.6.7). Trimmed reads were aligned using STAR (version 2.7.8a), with flags--chimSegmentMin 12, --chimJunctionOverhangMin 8, --chimOutJunctionFormat 1 to detect gene fusions, and gene fusions were quantified using STAR-fusion (version 1.10.0).^80^ Trimmed reads were also used to quantify transcript-level abundance using Kallisto (version 0.46.2) and transcripts were annotated using gencode version 23.^93^

### Neoantigen prediction

Variant annotation: Each variant in the final variant set was HLA-typed using OpiType (version 1.3.5).^94^ The variant effect predictor (vep, version 103) from ENSEMBL was then used to annotate somatic mutations compared to the reference genome, hg19.^95^

Neoantigen prediction: For all missense, frameshift or in-frame variants, PVACseq from PVACtools (version 2.0.1) was used to predict neoantigens.^96^ Binding affinity of each peptide to MHC was predicted by the MHCFlurry algorithm and is represented by an IC50 score (high binding affinity is called at IC50 > 500).^97^ Neoantigens were further annotated by gene, variant type, protein change, HLA allele, mutant and wild type epitope sequence, RNA and DNA coverage of the variant, as well as prevalence of the mutation from OncoPhase.^90^

### Shared mutations

Mutation overlap: Mutations were called for both the DigiPico and FT samples compared to germline PBMCs. Each mutation was assigned a unique mutation ID (<CHR>-<POS>--<ALT>) and the intersection of mutation IDs for each sample was computed. Of the 14,767 called in the DigiPico sample and the 13,724 called in the FT, 6 mutations were present in both. These 6 mutations were checked in IGV to verify that they were confidently in both samples and absent from the germline sample, and 5/6 of these were validated like so. Mutation prevalence: Prevalence was calculated for each of these mutations using Oncophase (v 0.1).^90^

## QUANTIFICATION AND STATISTICAL ANALYSIS

Data was analyzed using GraphPad Prism version 10.4.1. Group sizes and the definition of error bars are indicated in figure legends. In some indicated bar graphs, the background is subtracted from the signal, and negative values are set to zero. Statistical analysis was performed using a two-tailed Student’s t test. P values <0.05 were considered significant; significance values are indicated as * (*P* < 0.05), ** (*P* < 0.01), and *** (*P* < 0.003).

