## Supplementary Figures for "Single cell functional immunogenomics of the fallopian tube reveals a precursor immune surveillance network for ovarian cancer prevention"

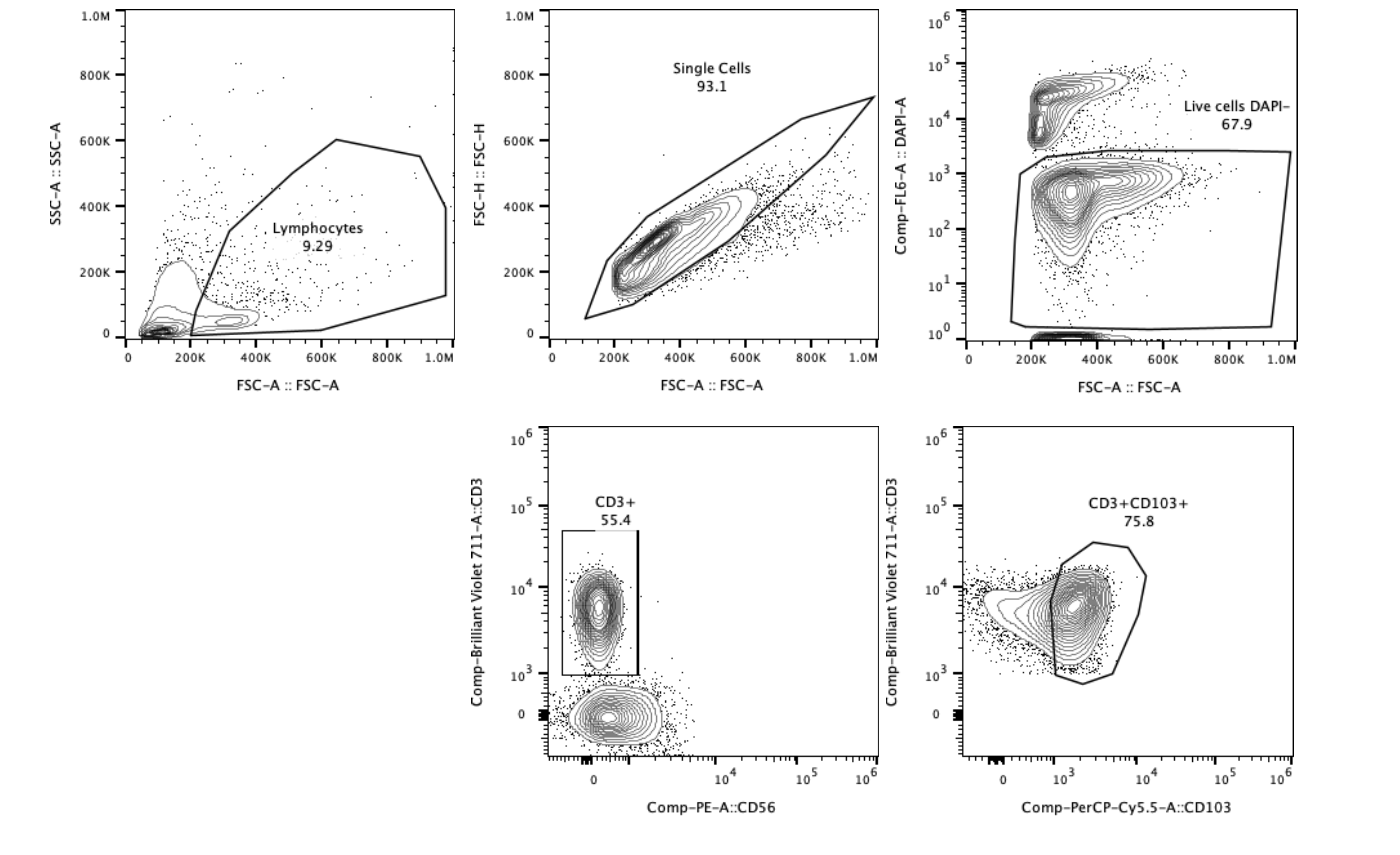

Figure S1 Gating strategy of tumor/normal FT/PBMC. Example of a normal FT.

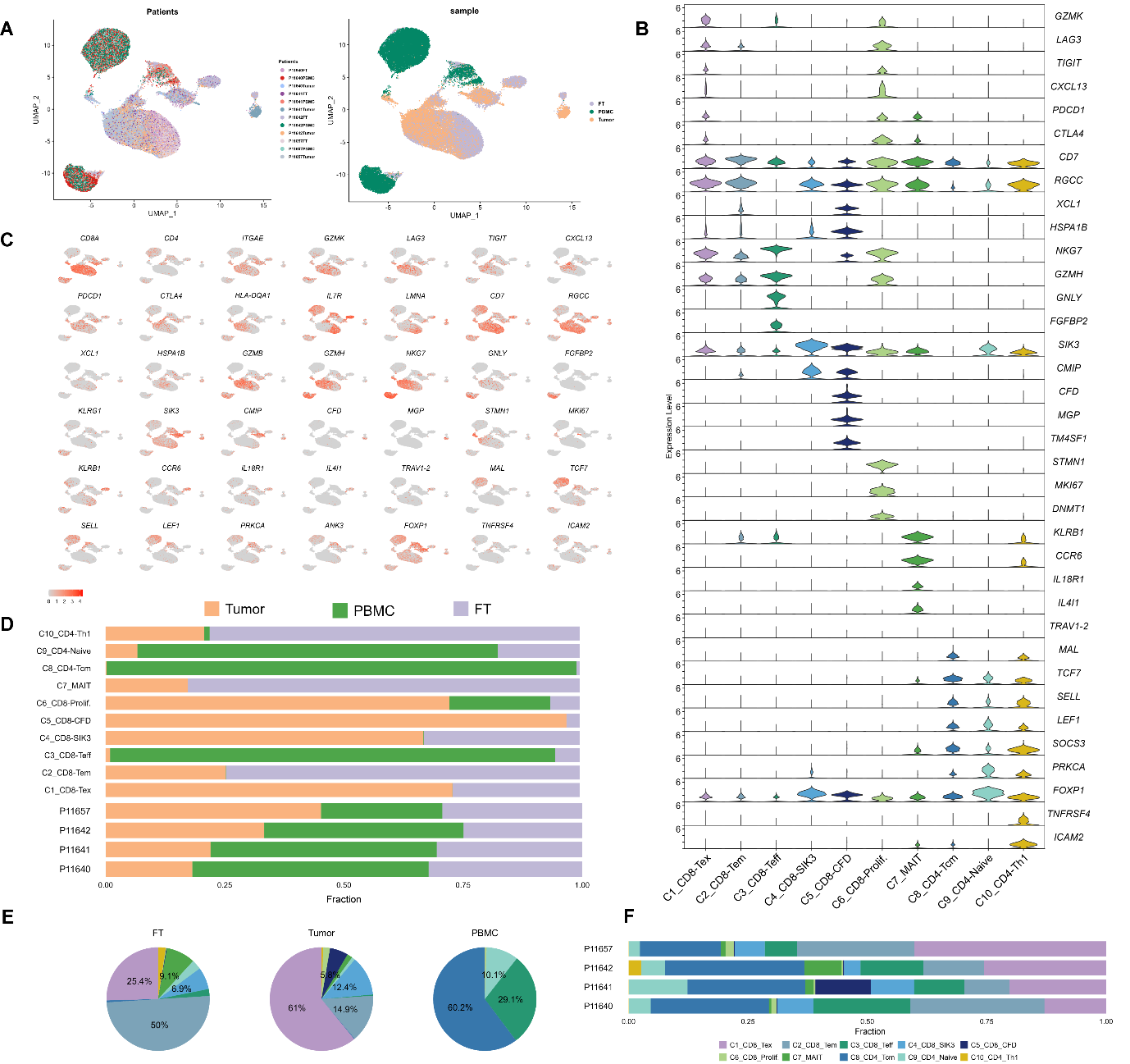

Figure S2 Characterization of T cell clusters and their dynamics in HGSOC.

The UMAP plot (A) displays T cells colored by patient origin on the left and by tissue site on the right. The violin plot (B) highlights the expression of canonical T cell subset marker genes, cell subset-specific genes, and key T cell checkpoint genes across the ten identified T cell clusters, with rows representing clusters and columns corresponding to signature genes. The UMAP red-scale projection (C) visualizes the expression levels of these marker and checkpoint genes across different clusters. The bar plots (D) depict the distribution of T cells from each tissue site across clusters at the top, while the bottom section shows the distribution of T cells by patient. The pie chart (E) presents the proportion of T cells within each cluster, categorized by tissue type, including FT, PBMC, and tumor samples. Lastly, bar plots (F) illustrate the cluster distribution of T cells across individual patients, providing insights into inter-patient variability in T cell composition within the HGSOC microenvironment.

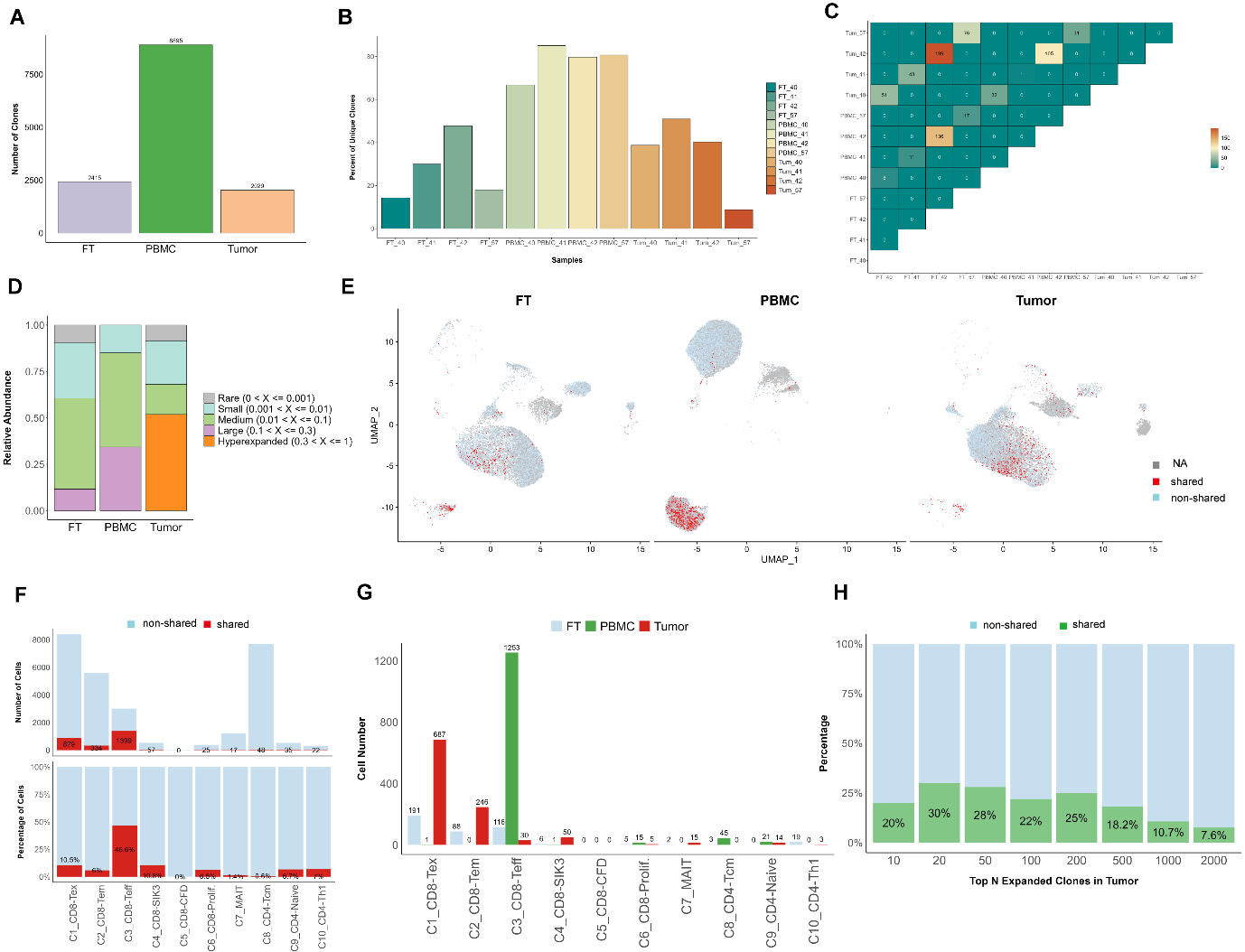

Figure S3 Detailed characterization of T cells with shared TCRs between PBMC and tumor tissue in HGSOC patients.

The bar plot (A) illustrates the number of unique TCR clones identified across different tissue sites. The bar plots in (B) depict the distribution of T cell clone sizes in each patient, stratified by tissue site. The clonal overlap plot (C) quantifies the number of shared TCRs between PBMC and tumor for each patient. Clonal abundance across tissue sites is shown in (D), where T cell clones are categorized based on expansion levels: rare (0 < X ≤ 0.001), small (0.001 < X ≤ 0.01), medium (0.01 < X ≤ 0.1), large (0.1 < X ≤ 0.3), and hyperexpanded (0.3 < X ≤ 1), revealing the expansion dynamics of shared clonotypes. The UMAP plot (E) highlights T cells with shared TCRs between tumor and PBMC (red), while non-shared TCRs are marked in light blue, and T cells lacking detected TCRs are in grey. The bar plots (F) display the number and proportion of shared T cells between PBMC and tumor across different clusters. In (G), the shared TCRs across tumor, PBMC, and FT are visualized, showing the number of shared T cells in each cluster and tissue site (FT in blue, PBMC in green, tumor in red). The bar plot (H) presents the proportion of shared TCRs between PBMC and the top N expanded clones in tumors, ranging from the top 10 to top 2000 most expanded clones. These analyses provide insights into the immune dynamics of PBMC-derived T cells and their potential interactions with the tumor microenvironment.

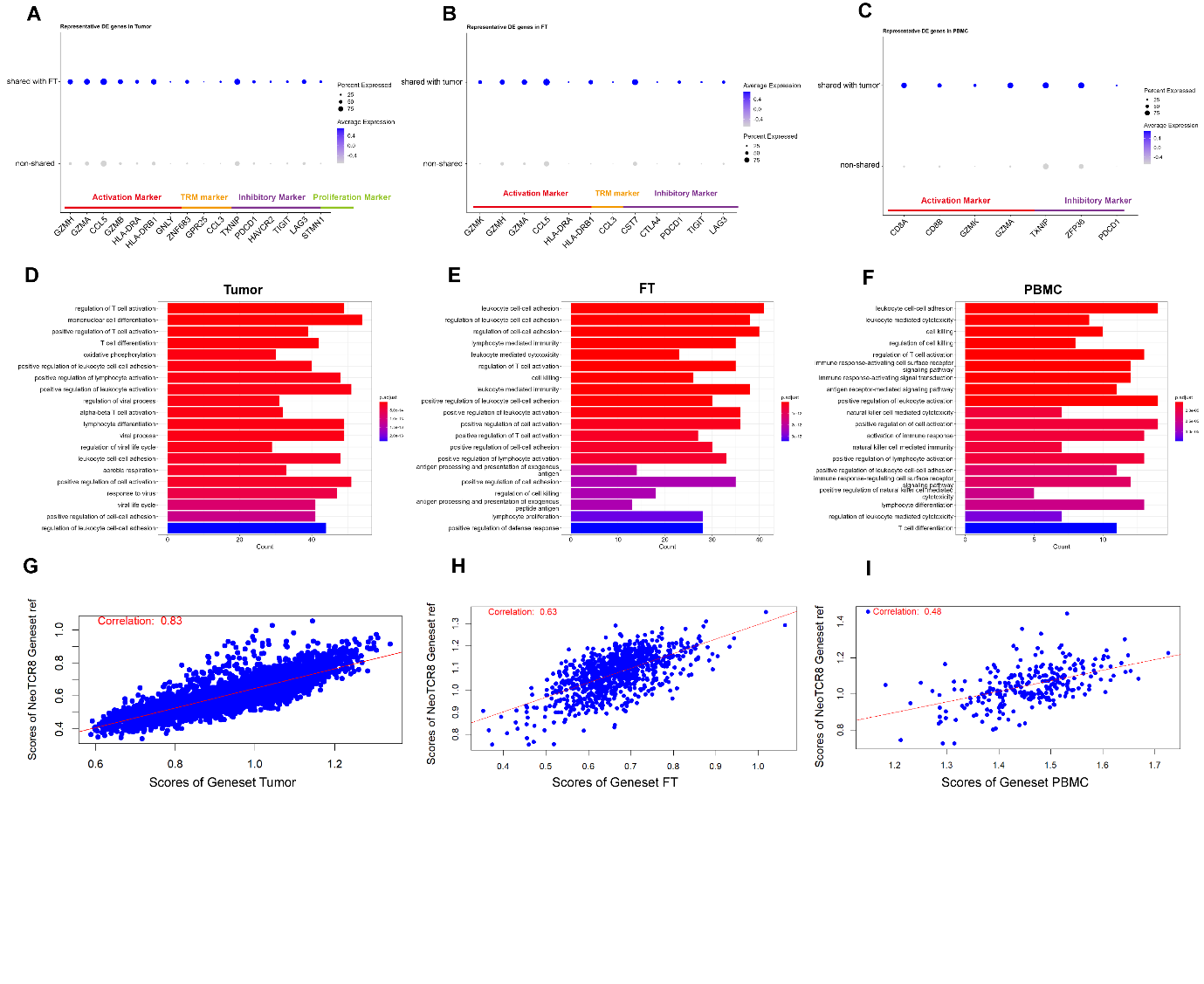

Figure S4 The transcriptomic differences between T cells with shared TCRs in tumors and non-shared TCRs across different tissue sites.

(A-C) Dot plots illustrate representative differentially expressed (DE) genes between shared and non-shared TCRs in tumor(A), FT (B), and PBMC (C).

(D-F) Gene Ontology (GO) enrichment analyses highlight pathways associated with differentially expressed genes in tumor (D), FT (E), and PBMC (F).

(G-I) Correlation plots assess the relationship between DE gene sets from shared versus non-shared TCRs and a published geneset of neoantigen-reactive CD8 T cells (NeoTCR8 Geneset ref.), analyzed separately in tumor (G), FT (H), and PBMC (I) datasets.

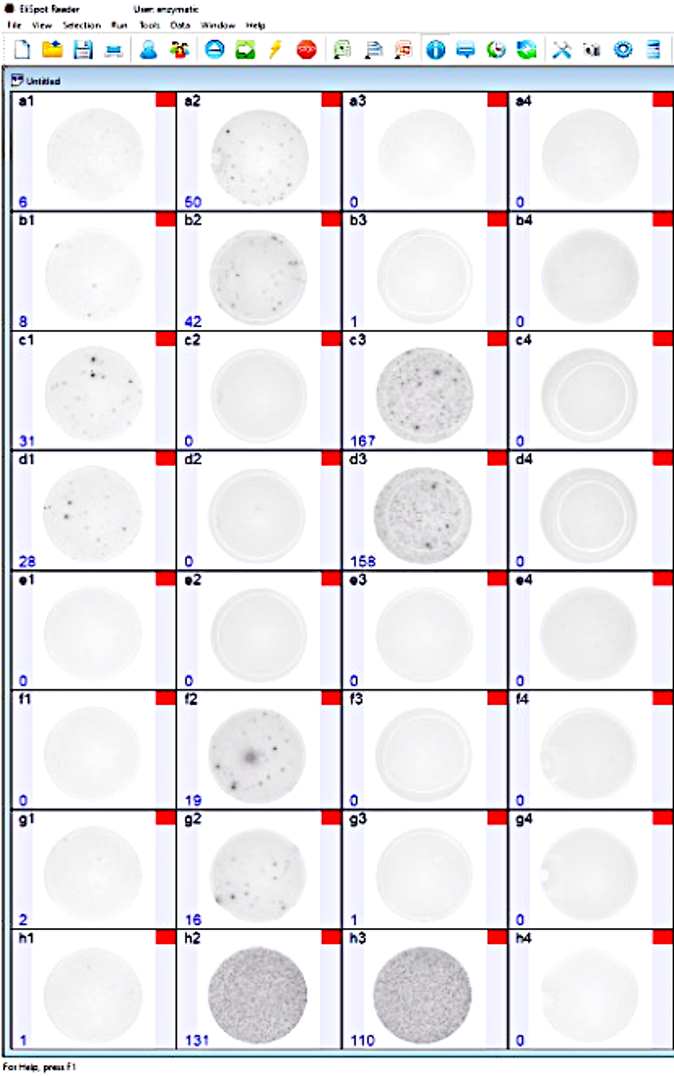

Figure S5 A representative image of interferon-γ (IFN-γ) ELISPOT wells, illustrating the immune response of TRMLs from FT upon ex vivo stimulation. Columns 1 and 2 display the ELISPOT results for TRMLs from the FT of patient 11642 following stimulation with medium alone (n=4; a1-b1, a4-b4), tumor organoids (n=4; c1-d1 with 1.5×10⁴ TRMLs and a2-b2 with 2×10⁵ TRMLs), or FT organoids (n=4; e1, f1, g1, and h1). FT TRMs stimulated with TransAct nanobeads served as a positive assay control (h2). Column 3 shows ELISPOT wells of TRMLs from patient 11639, either unstimulated (a3-b3) or co-cultured with tumor organoids derived from the patient’s FT (c3-d3), eliciting a strong IFN-γ response. However, due to the difficulty of identifying normal FT tissue from this patient to establish FT organoids as a control, further comparative analysis was not performed.

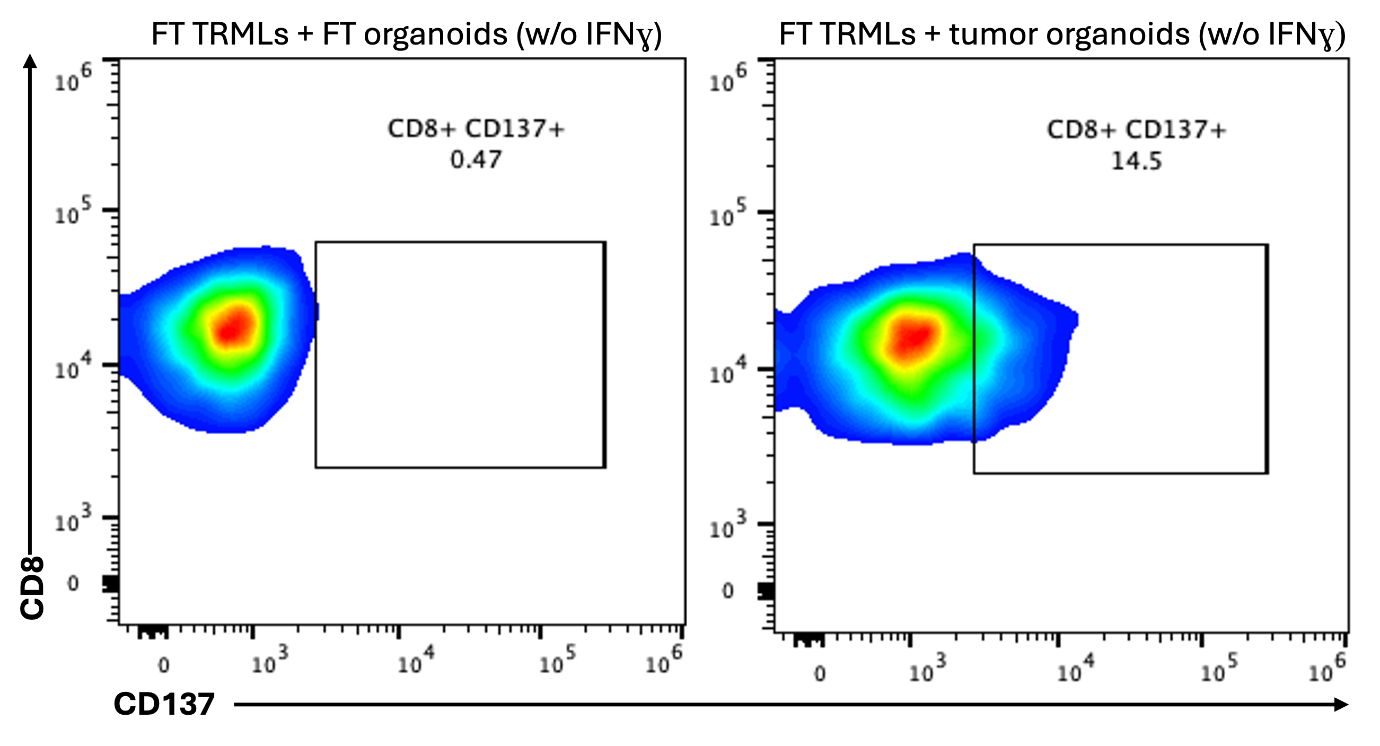

Figure S6. Dot plots of CD3 and CD137 expression by tumour reactive tissue resident memory T cells (TRMLs) in FT, following coculture with or without tumour organoids (with no prior pre-stimulation with IFNɣ).

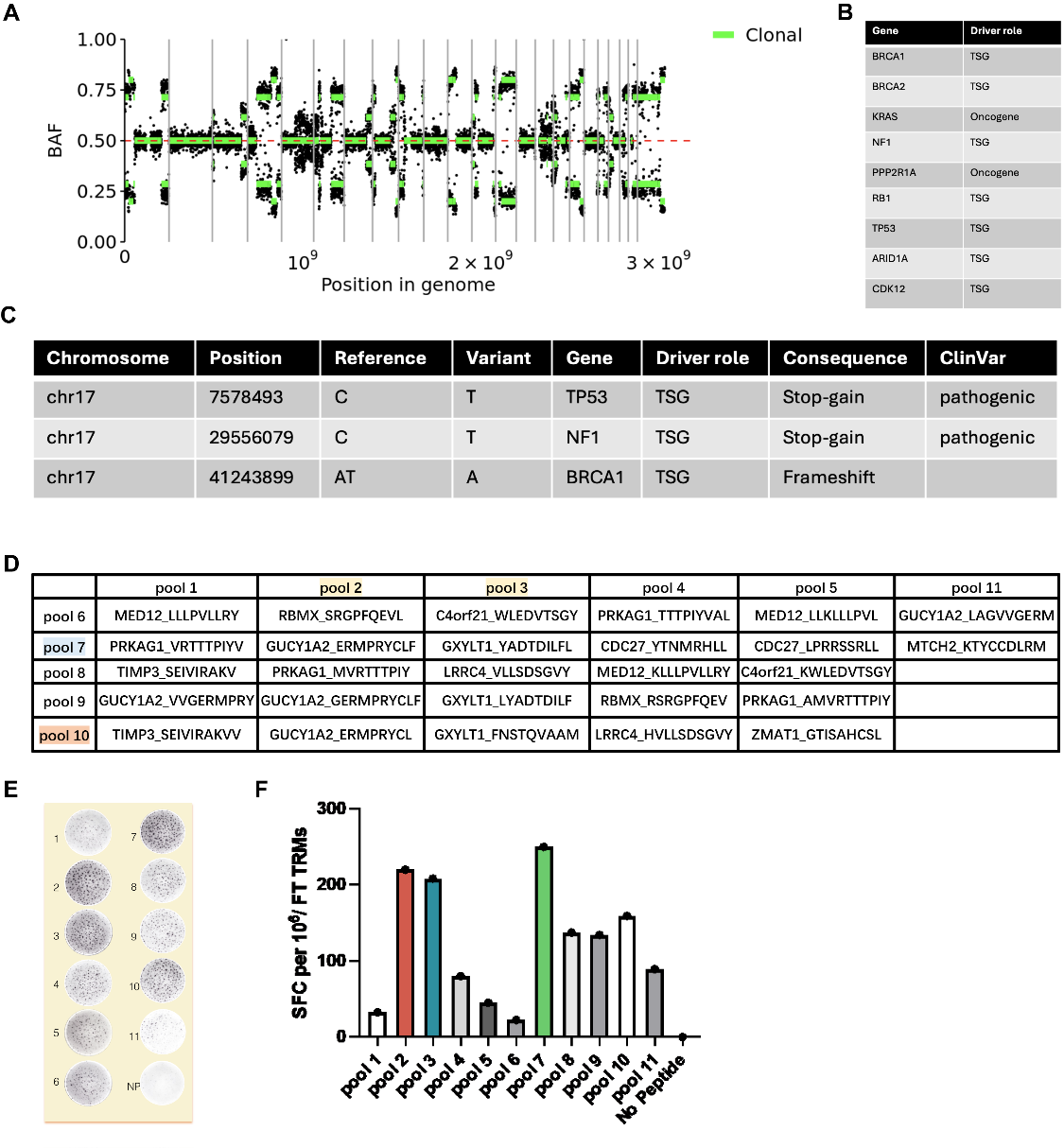

Figure S7 Whole-genome sequencing (WGS) analysis of the tumor, reveals key genomic features of high-grade serous ovarian cancer (HGSOC).

(A) B allele frequency (BAF) plot showing genome instability, highlighting profound chromosomal alterations characteristic of HGSOC.

(B) Copy number variations (CNVs) are illustrated, showing amplifications and deletions across the genome at the site of selected important oncogenes or tumor suppressor genes (TSGs).

(C) The presence of driver mutations is identified, providing insights into the molecular mechanisms underlying tumor progression.

(D) The peptide pools tested in (E) are detailed in a table listing their individual components.

(E) Representative ELISA spots display IFN-γ responses to eleven pools of 9-mer peptides, indicating immune recognition of potential neoantigens.

(F) The number of IFN-γ spot-forming cells (SFCs) per 10⁶ FT TRMs is quantified, demonstrating the extent of immune activation.
